# Glucose limitation in *Lactococcus* shapes a single-peaked fitness landscape exposing membrane occupancy as a constraint

**DOI:** 10.1101/147926

**Authors:** Claire E. Price, Filipe Branco dos Santos, Anne Hesseling, Jaakko J. Uusitalo, Herwig Bachmann, Vera Benavente, Anisha Goel, Jan Berkhout, Frank J. Bruggeman, Siewert-Jan Marrink, Manolo Montalban-Lopez, Anne de Jong, Jan Kok, Douwe Molenaar, Bert Poolman, Bas Teusink, Oscar P. Kuipers

## Abstract

A central theme in biology is to understand the molecular basis of fitness: which strategies succeed under which conditions; how are they mechanistically implemented; and which constraints shape trade-offs between alternative strategies. We approached these questions with parallel bacterial evolution experiments in chemostats. Chemostats provide a constant environment with a defined resource limitation (glucose), in which the growth rate can be controlled. Using *Lactococcus lactis*, we found a single mutation in a global regulator of carbon metabolism, CcpA, to confer predictable fitness improvements across multiple growth rates. *In silico* protein structural analysis complemented with biochemical and phenotypic assays, show that the mutation reprograms the CcpA regulon, specifically targeting transporters. This supports that membrane occupancy, rather than biosynthetic capacity, is the dominant constraint for the observed fitness enhancement. It also demonstrates that cells can modulate a pleiotropic regulator to work around limiting constraints.

## INTRODUCTION

Many microorganisms have a remarkable versatility to adapt to different environments, either at the short time scale through signaling and gene expression networks, or at an evolutionary time scale through genomic adaptations to selective pressures. A key question in (micro)biology, is how specific selective pressures shape molecular strategies, how evolvable they are, and which constraints and resulting trade-offs in phenotypic traits are relevant under what condition.

In recent years a number of studies (Dekel and Alon, 2005; Molenaar *et al.*, 2009; Scott *et al.*, 2010, 2014) pointed towards cellular resource limitations, born from environmental (nutritional) constraints but also from intrinsic kinetic and physicochemical constraints, as main forces for regulatory strategies. These relate to the role of protein costs in particular, and how the need for ribosomal protein synthesis capacity requires growth-rate dependent regulation of the proteome (Scott *et al.*, 2010). Although these concepts can explain many regulatory phenomena, such as catabolite repression (Görke and Stülke, 2008) and overflow metabolism (Basan *et al.*, 2015), it remains to be demonstrated (i) to what extent such strategies require optimality or are optimal solutions to the resource allocation problem; (ii) what exactly the constraints are that govern the resource allocation; (iii) what the regulatory mechanisms are that result in proteome re-allocation; and (iv) whether these concepts apply to microorganisms other than *E. coli* – the organism for which these concepts have been largely developed and experimentally tested.

We have recently challenged the view that protein costs play a role in metabolic switches in the lactic acid bacterium *Lactococcus lactis*: we found a switch from mixed acid fermentation to homolactic fermentation (an anaerobic variant of the Crabtree effect) in the absence of proteome alterations (Goel, Eckhardt, Puri, Jong, Branco dos Santos, Giera, Fusetti, Vos, Kok, Poolman, Molenaar, Kuipers and Teusink, 2015). One explanation was that this bacterium might not display optimal behaviour in the chemostat in which they were cultivated. This primed us to do laboratory evolution experiments as they offer unique opportunities for the testing of adaptive strategies, whilst monitoring evolving populations under well-defined conditions.

Laboratory evolution of microorganisms can be carried out under a broad range of selective pressures, such as varying growth rate in serial batch cultures (Bachmann *et al.*, 2012; Wiser, Ribeck and Lenski, 2013), antibiotic resistance in different cultivation settings (Lázár *et al.*, 2013), growth rate under nutrient limitation in chemostats (Gresham and Hong, 2014), cell number in emulsion-based systems (Bachmann *et al.*, 2013), and growth rate in dynamic environments (Meadows *et al.*, 2010). However, to study the role of limited resource allocation on adaptive strategies, at different growth rates and evolutionary time scales, the chemostat is perfect (Gresham and Hong, 2014). It allows for full control over the nature of the resource limitation and the growth rate, and provide a constant, steady-state environment. We studied microbial evolution of *Lactococcus lactis* in highly-standardized glucose-limited chemostats at two different dilution rates with four replicates each. We found the exact same residue of a pleiotropic regulator to be mutated in all of these experiments, and confirm the fitness-enhancing effect of this mutation in chemostat competition experiments at other dilution rates as well. Detailed characterization of the biochemical and phenotypical consequences of the mutation demonstrated a reprogramming of the regulatory logic of the regulator, by inducing the suppression of alternative carbon uptake systems under low glucose levels whilst maintaining the –otherwise similarly regulated- high affinity glucose transporter. The occurrence of one dominant mutation suggests a single-peaked fitness landscape and shows that the dominant constraint under such chemostat conditions is not cytosolic protein synthesis costs, but rather membrane occupancy. We thus experimentally confirm and extend previous computational studies in *E coli* (O’Brien *et al.*, 2013) which suggested the action of different constraints (nutrient limitation and proteome limitation) under different conditions. While doing so, we provide a full mechanistic mapping from mutation to regulation to fitness.

## RESULTS

### Chemostat and cultivation design for long-term microbial evolution under stable environmental conditions

Until now, microbial evolution in chemostats has been rather limited by the number of replicates and by the number of generations, *i.e.* volume changes. Initially described as a device that could continuously cultivate bacteria for an indefinite period of time (Novick and Szilard, 1950), to date the longest published cultivation using a chemostat is only a few hundred generations (Gresham and Hong, 2014). This is several orders of magnitude shorter than the longest serial batch cultivation experiments – 50,000 generations and counting (Wiser, Ribeck and Lenski, 2013). To address the failure of long-term continuous cultivation, we developed a versatile bioreactor cap and experimental set-up to perform parallel, prolonged cultivations under chemostat conditions, at small working volumes (detailed in Supplementary Materials and Figure 1). These advances allowed us to conduct experiments in quadruplicate for more than 1000 consecutive generations.

**Figure 1.**
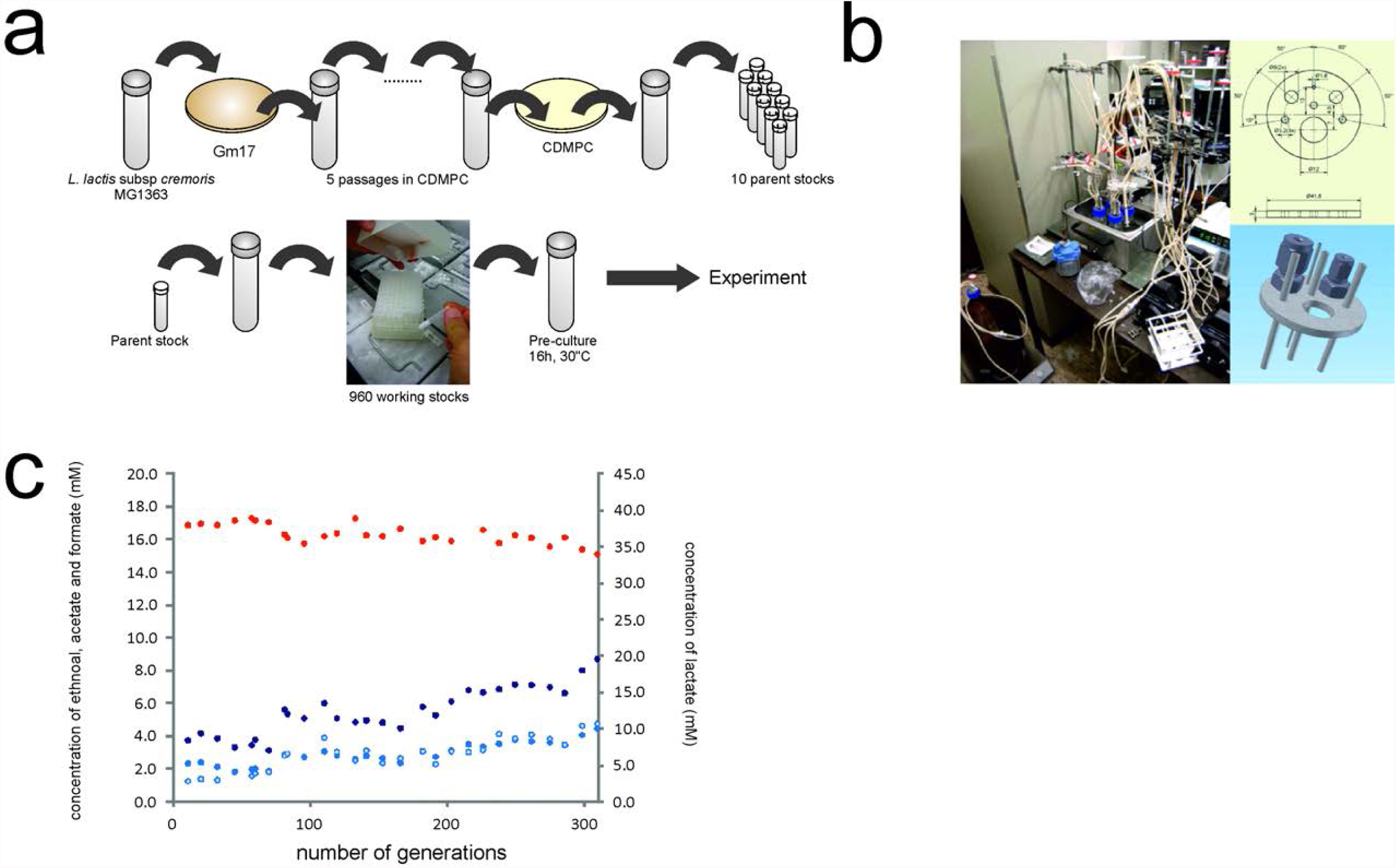
(a) Standardized cryopreservation procedure to minimize strain variability among experiments. GM17: M17 medium supplemented with 25 mM glucose, CDMPC: chemically defined medium for prolonged cultivation. (b) Miniaturized chemostat set up to allow prolonged cultivation experiments in parallel and (c) the online measurements of organic acids for strain 309C1 taken during the evolution experiments at D of 0.5 h-1 showing lactate (red), formate (dark blue), acetone (blue) and ethanol (light blue).

Additionally, a requirement of laboratory evolution is that sources of technical variability between replicates are minimized. Our initial focus was set on carefully defining the cultivation conditions (chemostat and medium) and on standardizing experimental procedures (inoculation, sampling, and cell storage). We developed a chemically defined medium particularly suited for prolonged cultivation (CDMPC) based on the nutrient requirements (Jensen and Hammer, 1993) and biomass composition of *L. lactis* MG1363 (Oliveira, Nielsen and Förster, 2005) (detailed in supplementary materials). To reduce inoculum variability we implemented a strictly controlled cryopreservation and inoculation procedure (Figure 1 and detailed in Supplementary Materials).

The cultivation history of bacterial strains is of upmost importance. Strains can rapidly deviate from their original genotype, which can lead to noticeable phenotypic differences (Linares, Kok and Poolman, 2010; Bachmann *et al.*, 2012). Our initial inoculum came from the original *L. lactis* MG1363 sequenced stock (Linares, Kok and Poolman, 2010). Since our newly designed medium could potentially lead to the selection of new genotypes, we performed a short-term serial batch adaptation of this strain to the medium CDMPC. The resulting single isolate picked (Genr0) was cryopreserved and used as the seeding population for all the laboratory evolution experiments performed.

### Parallel laboratory evolution in chemostat cultures

We exposed *L. lactis* MG1363 to parallel chemostat cultivations: Genr0 was cultivated continuously in quadruplicate, each using 60 mL glucose-limited chemostats at a growth rate of 0.5h^-1^ for over 300 volume changes. Culture samples were harvested periodically every 10-15 generations from the effluent to determine cell density and the fluxes of fermentation substrates and products (an example is given in Figure 1C). Cell samples were stored as glycerol stocks at -80°C to be used as snapshots of the evolutionary process.

Throughout the prolonged cultivation, all four replicates (referred hereon as 309C1 through 309C4) displayed very similar behavior. Biomass concentration did not vary significantly indicating that biomass yield on substrate remained fairly constant (Supplementary Figure 1). *L. lactis* can catabolize glucose through homolactic or mixed acid fermentation, excreting, respectively, lactate, or acetate, formate and ethanol (Thomas, Ellwood and Longyear, 1979). At a dilution rate of 0.5 h^-1^, Genr0 produces mainly lactate (lactate:acetate ratio ~16). Throughout the course of the experiment, all parallel reactors shifted gradually towards mixed acid fermentation leading to a ratio of approximately 7. Despite the fact that this fermentative mode leads to a theoretical 50% increase in ATP yield, the biomass concentration did not change. The latter can be explained by the observation that the evolved cells also gradually excreted more pyruvate, the final shared precursor of the homolactic and the mixed acid fermentative pathways (Supplementary Figure 1).

When evolved cells were grown on glucose in batch, we observed differences with Genr0 in the maximal growth rate (μ_max)_, length of the lag phase and sedimentation. We compared to Genr0 both the population samples collected after 309 generations and single colony isolates from the evolved population samples (Supplementary Figures 1 and 6). Irrespective of whether the evolved populations or the isolates were compared, they were clearly outperformed by Genr0 in batch. Most notably, μ_max_ dropped to a value close to the dilution rate at which cells were evolved, which falls nearly 25% below the μ_max_ of the parent strain. The evolved strains also exhibited extended lag phases and sedimentation during batch cultivations. The sedimentation was reminiscent of the AcmA-deficient phenotype previously described (Buist *et al.*, 1995).

### Identification of the mutations arising during the evolutionary experiment

We sequenced single-colony isolates from the original strain stock of Genr0 and from the end of the four replicate evolution experiments performed at a dilution rate of 0.5h^-1^ (*i.e.* 309C1, 309C2, 309C3, 309C4). The published whole genome sequence for *L. lactis* MG1363 (Linares, Kok and Poolman, 2010) was used as a reference for sequence assembly. Mutations were identified and verified by Sanger sequencing (Supplementary Table 2). In comparison with Genr0, we found SNPs unique to the evolved strains (Figure 2). The number of SNPs accumulated per genome was between one (309C1) and six (309C4). This corresponds to a mutation rate of 1.3 to 7.8 x 10^-9^ (per base pair per generation) and is in line with currently published estimates (Barrick *et al.*, 2009).

**Figure 2.**
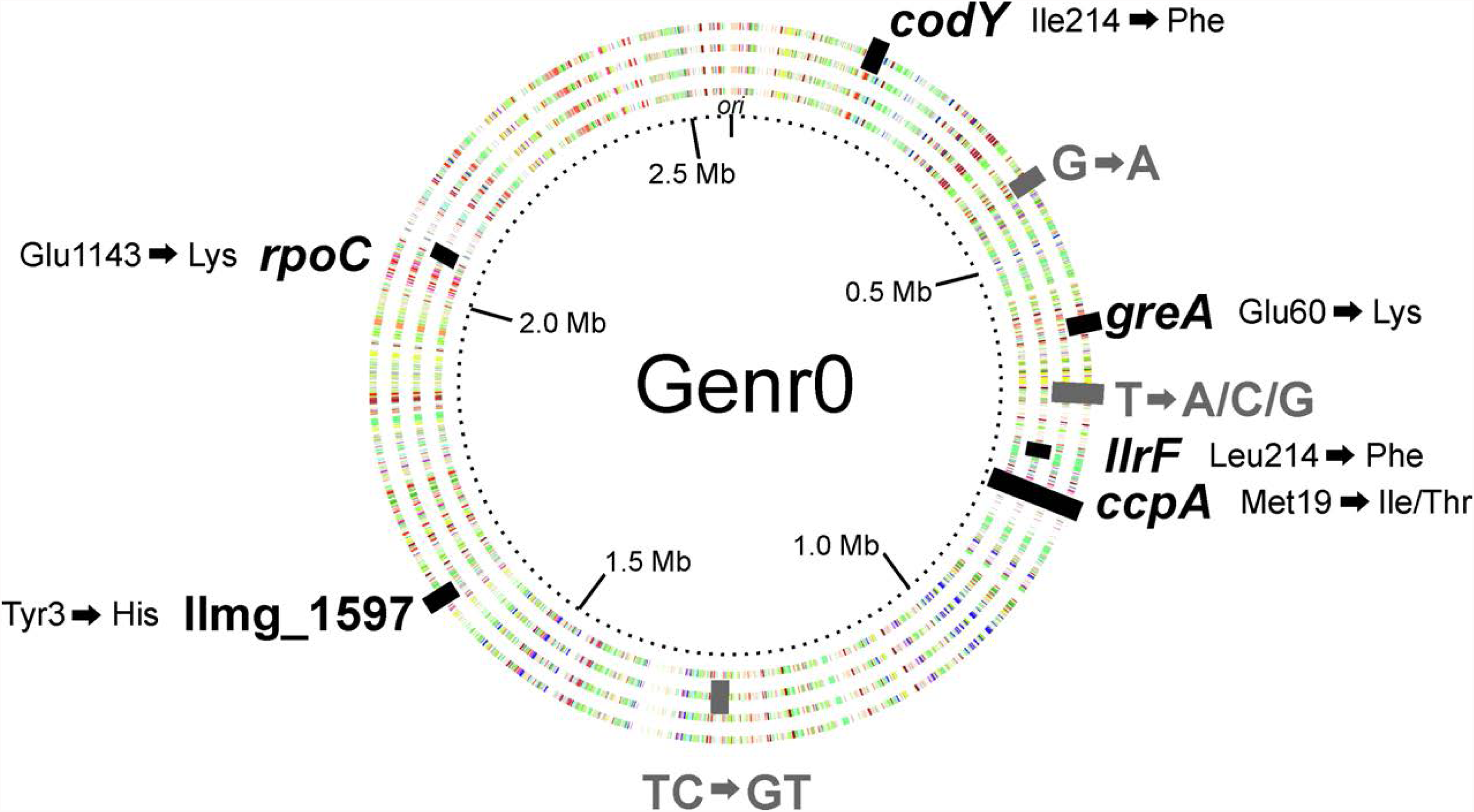
Mutations identified in the 4 evolved strains compared to the original strain Genr0. The genes and intergenic regions with SNP mutations are shown. For coding regions (in black) the resulting amino acid substitution is indicated. The dotted inner ring indicates the genome position. The rings are color-coded according to the COG-classification of the gene present on the forward and reverse strands, and from inner to outer ring we represent evolved strains 309C1 to C4.

All of the SNPs in the coding regions were non-synonymous (more details in Supplementary Materials), and strikingly, all strains accumulated a mutation in the codon coding for Met19 in the global transcriptional regulator CcpA of Genr0. At the DNA level, the mutations differed between the evolved strains. In strains 309C1, 309C2, and 309C3, ATG was mutated to either ATA or ATC, resulting in the amino acid change from Met to Ile. In none of the sequencing experiments performed for cells cultured at a D of 0.5h^-1^ was the Ile codon ATT identified (see also section on *Dynamics of Evolution Experiments*). For strain 309C4 the ATG codon was mutated to ACG resulting in a Met to Thr change. Strains 309C2, 309C3 and 309C4 contain a few additional mutations, while the isolate of 309C1 only contains the SNP in *ccpA*. Since 309C1 exhibits very similar phenotypes to those of the other strains, this mutation must be sufficient to confer a growth advantage to the evolved strains when compared to Genr0, and to a great extent be the genetic basis of the common phenotypic differences observed. These include the growth kinetics and sedimentation phenotype (Supplementary Figures 1 and 6) and, as addressed below, altered glucose utilization kinetics. The sequences were also analyzed for insertions and/or deletions and when compared to the Genr0 sequence; the evolved strains showed no major frameshifts (see Supplementary Table 2).

### Effect of CcpA M19I on global gene expression patterns

The evolved strains were revived in CDMPC and grown until mid-exponential phase at which point microarray analyses were performed (Kuipers *et al.*, 2002). Gene expression in the evolved strains was compared to that of Genr0. Evolved strain 309C1, which contains only the SNP in *ccpA,* showed an altered gene expression dominated by genes involved in membrane transport, especially carbohydrate import (Supplementary Figure 3, Supplementary Table 3, Supplementary Table 4). CcpA controls the preferential use of glucose over other sugars (Deutscher, Francke and Postma, 2006) and regulon analysis of the differentially expressed genes revealed that CcpA-regulated genes were indeed over-represented in the data set.

*L. lactis* contains three glucose import systems – two phosphotransferase systems, PTS^Man^ and PTS^Cel^, and a glucose facilitated diffusion system (Castro *et al.*, 2009) – and expression of the genes encoding the high affinity PTS^Man^ were up-regulated compared to the original strain, while those encoding the low affinity PTS^Cel^ were down-regulated (Figure 3A). The effects of these gene expression changes were investigated further by monitoring the uptake of ^14^C-glucose in whole cells. In accordance with the sequencing and gene expression data, *i.e.* the glucose transporters themselves are not mutated but rather their expression is changed: the rate at which glucose was taken up by the cells was increased nearly 3-fold in the evolved strain, while the apparent Michaelis constant for glucose was unchanged (Figure 3B).

**Figure 3.**
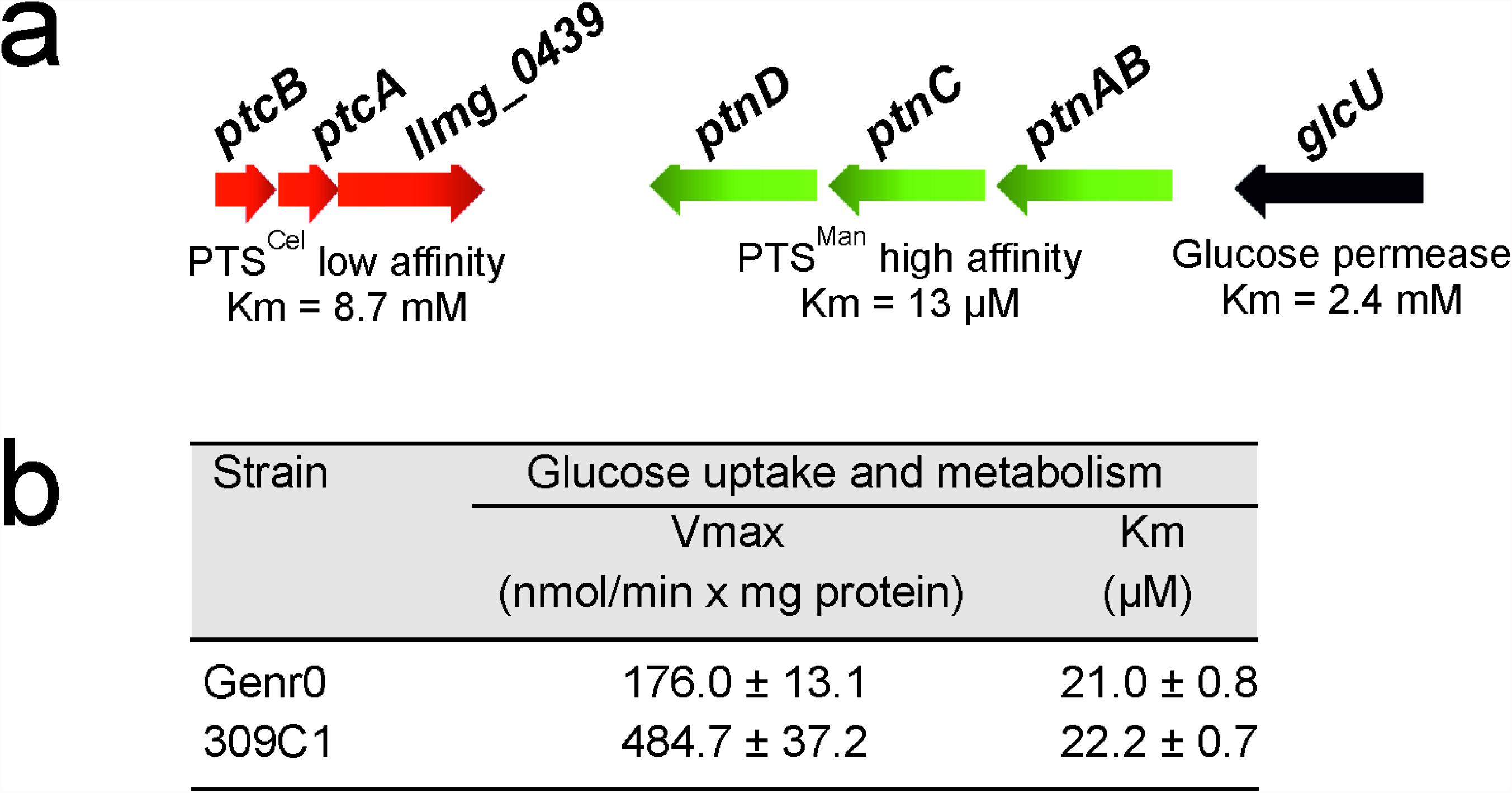
Altered gene expression in strain 309C1 is focused on transport pathways. (a) The expression of genes involved in glucose uptake in *L. lactis* MG1363 was significantly changed in the evolved strains. Highlighted in green are genes that were up-regulated and in red those that were down-regulated. (b) Kinetic parameters of glucose transport in Genr0 and strain 309C1 were determined in cells grown to exponential phase in CDMPC. Glucose transport was assayed with [^14^-C]-labeled glucose. Values of three independent experiments were averaged and are reported ±SD. V_max_ and K_m_ were determined using glucose concentrations from 1.2 to 200 μM (Gouridis *et al.*, 2015).

The ability to utilize other carbon sources was diminished for the evolved strains (Supplementary Figure 4A). This coincided with the down regulation of, amongst others, genes encoding for maltose transporters, ABC sugar importers and sugar utilization enzymes (Supplementary Table 3), many of which are CcpA-regulated.

In general, CcpA is a versatile global regulator that can act as a transcriptional repressor or activator. An amino acid substitution in the DNA binding region may therefore have qualitatively different effects at different target promoters. This is indeed what we observe: 15 CcpA-regulated genes are down regulated, 6 are up-regulated, while 11 remain unchanged. We next examined the molecular basis of the differential effects on expression.

### Effect of M19I mutation on the binding affinity of CcpA

Met19 is located in the second helix of the DNA-binding domain and is highly conserved in CcpA (Figure 4A and Supplementary Figure 8). *L. lactis* CcpA binds to *cre* sites for which the consensus sequence is known: TGNNANCGNTTNCA. Sequence analysis revealed that putative *cre* sites upstream down-regulated genes were closest to the *cre* consensus sequence, while up-regulated genes deviated and were less likely to have both C7, G8 and C13 but always T10 (Figure 4B). We tested this further by analyzing CcpA with either Met or Ile or Thr at position 19 and a select number of *cre* site sequences in *in vitro* binding assays and molecular dynamic simulations.

**Figure 4.**
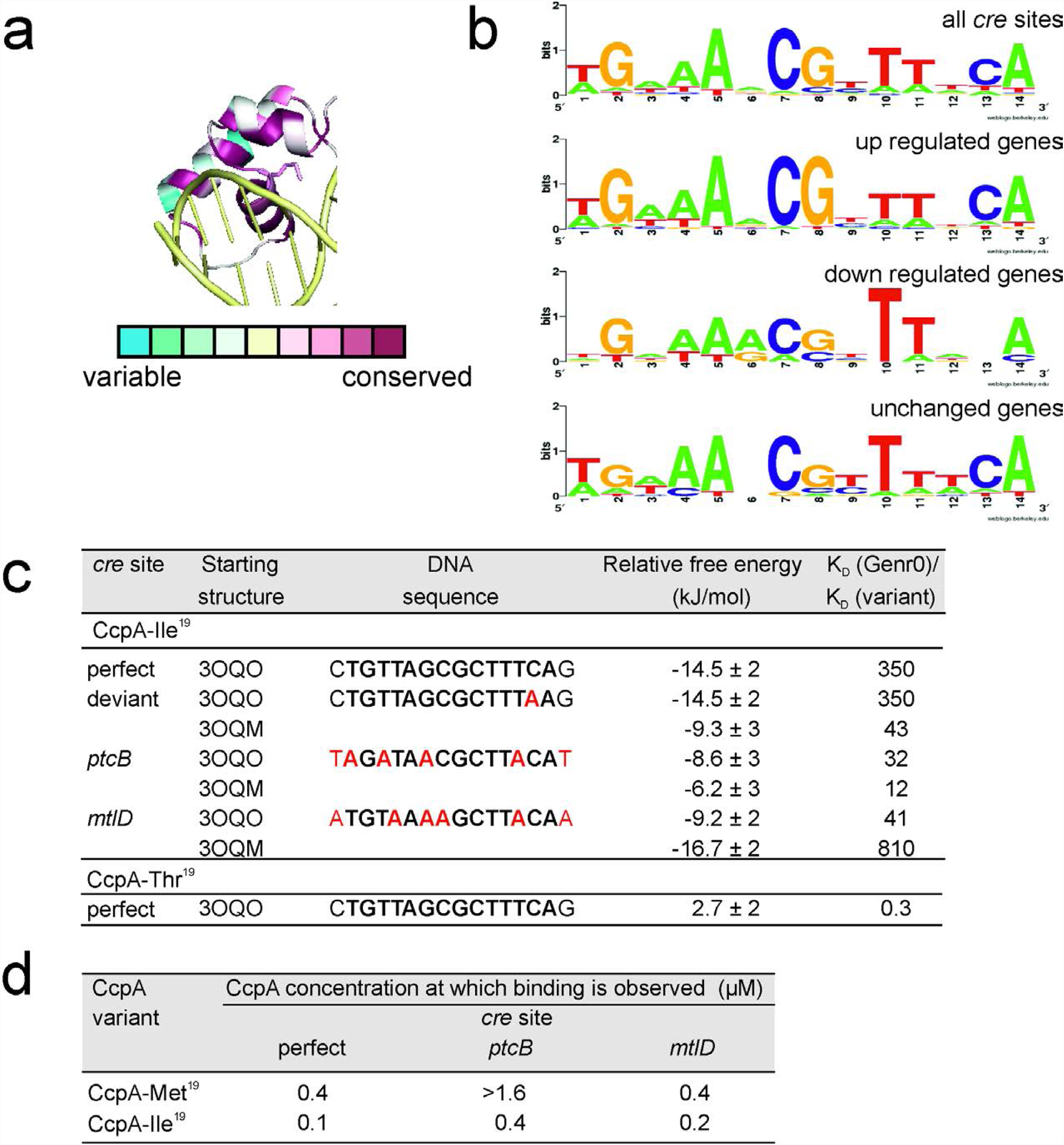
Mutations in *ccpA* result in increased binding affinity for certain *cre* sites. Mutations in the Met^19^ codon in the DNA binding domain of CcpA alter the binding affinity of CcpA for certain *cre* sites. (a) View of the DNA-binding domain with the side chain of Met19. Using ConSurf 2010 (Ashkenazy *et al.*, 2010) and the *B. subtilis* CcpA-HPr complex bound to a synthetic cre site (3OQN) as a starting structure, the level of amino acid sequence conservation amongst LacI transcriptional regulators was analyzed. CcpA is colored according to the level of conservation within the LacI family of transcriptional regulators, while the DNA is colored yellow. Dark purple indicates 100% conservation, while blue indicates extensive sequence variation. (b) Sequence logo diagrams representing the abundance and position of nucleotides in the CcpA-regulated genes of *L. lactis* MG1363. (c) Changes in relative binding free energy upon substitutions at Met19 in CcpA as calculated by molecular dynamics. The reported errors of the relative free energies are standard errors of the mean. (d) Binding of CcpA to *cre* sites *in vitro*. The binding of CcpA-Met^19^ and CppA-Ile^19^ was tested with DNA sequences identified as *cre* sites upstream of *ptcB* and *mtlD* as well as a perfect *cre* site. As a negative control the CodY recognition site upstream of *oppD* was also tested (See also Supplementary Figure 10).

For all 4 *cre* sites tested *in vitro*, the CcpA-Ile19 variant showed an increase in binding affinity for the DNA operators selected (Figure 4D and Supplementary Figure 10). This was most pronounced for the sequence upstream of *ptcB* and least pronounced for *mtlD*. To complement the binding assays, molecular dynamic simulations were performed. This allowed us to determine the relative binding free energy of the CcpA variants to a similar set of DNA operators (Figure 4C, Supplementary Figure 9).

The effect of either the Ile or Thr substitution at position 19 was first investigated by calculating the relative binding free energies of CcpA to a canonical *cre* site (Schumacher *et al.*, 2011). The Ile variant had a more negative free energy of binding as compared to the wild type protein, whereas the Thr variant did not exhibit significantly different binding to the wild type protein. We then tested the effect of the DNA sequence on the binding free energy for wild type and evolved CcpA, using structures derived from CcpA bound to two different *cre* site sequences (3OQO and 3OQM) (Schumacher *et al.*, 2011). When the Ile was present at position 19, tighter binding to all DNA sequences tested was computed, irrespective of the starting structure used.

### Population dynamics of prolonged cultivation experiments

The frozen population stocks, collected periodically throughout the prolonged cultivations, served as snapshots of the adaptive evolution process to the chemostat environment. By determining the abundance of the CcpA-Ile19 variant in stocks collected at different generations, we were able to unravel the wash-in dynamics that outcompeted the native CcpA-Met19. Quantification was carried out using pyrosequencing of the region containing the SNP in the *ccpA* locus. The CcpA-Ile19 variant emerged after 50 generations in the evolved populations and became dominant already by generation 150 (Supplementary Figure 11). While Met is encoded by ATG, Ile can be encoded by ATA, ATC or ATT. However, only the first two codons were identified in the D=0.5 h^-1^ evolution experiment, with the ATA mutant emerging prior to ATC (Supplementary Figure 12). At generation 309, when the experiment was stopped, over 97% of the population carried the CcpA-Ile19 variant.

We then tested whether the phenotypic changes imposed by the CcpA mutation would be sufficient to explain the wash-in kinetics observed using the glycerol stocks. For this purpose, a simple model of chemostat growth with two competing strains was used to simulate the prolonged experiment at a D of 0.5h^-1^. The model was able to accurately fit the observed population dynamics, suggesting that the underlying mechanism can be fully understood in terms of chemostat wash-in kinetics and is attributable to the mutant CcpA regulator (Supplementary Figures 13A-C). Sensitivity analysis of this model showed that two pairs of parameters cannot be estimated independently, only their ratios (Supplementary Figures 13D-E). These are (i) the Monod constants (K_s_) of the parent (Genr0) and the evolved (309C1-C4) strains and (ii) the biomass yield of the parent strain and the emergence frequency of the fitter strain. This uncertainty has, however, no implications for the conclusions derived from the model. Another important aspect emerged from analyzing the model: the cells evolved at a D=0.5h^-1^ are predicted to outcompete Genr0 at any constant dilution rate not higher than μ_max_.

### Parallel laboratory evolution and competition experiments under chemostat conditions at different dilution rates

We next addressed whether the fitness effect of the CcpA-Ile19 variant is dependent on growth rate. We performed new evolution experiments in the chemostat environment at a dilution rate of 0.6h^-1^, since at a D closer to μ_max_ (~0.67 h^-1^), the concentration of limiting nutrient becomes higher (Goel, Eckhardt, Puri, Jong, Branco dos Santos, Giera, Fusetti, Vos, Kok, Poolman, Molenaar, Kuipers and Teusink, 2015), hence modulating the strength of the selection pressure applied. Experiments were carried out for as many as 800 volume changes, in a similar fashion as was done for the D of 0.5h^-1^. Sanger sequencing of the *ccpA* locus from fragments, obtained by PCR using the frozen stocks as templates, indicated that indeed the CcpA-Ile19 substitution was washing in as predicted by the model. Re-sequencing of the whole-genome of single isolates of each of the populations evolved at 0.6h^-1^ revealed that once again the only mutation that was consistent across all isolates was located in residue 19 of CcpA. This time all possible codons that encode Ile were found, suggesting that the amino acid substitution is the only factor that confers a strong fitness benefit.

We further tested whether the same mutation is also advantageous for dilution rates that are far lower than μ_max_ by performing competition experiments in chemostats between Genr0 and 309C1 (harboring only the CcpA-Met19 mutation). We used our growth model parameterized with the data from the evolution experiments at a D of 0.5h^-1^, to simulate the population dynamics of competition experiments carried out at a D of 0.2 and 0.3h^-1^ (Figure 5). Based on this, we designed and performed the experiments, in which a small fraction (~10%) of cells from a steady-state culture of 309C1 were mixed with a steady-state culture of Genr0 maintained at the same D. The population dynamics of these mixed cultures were monitored in time by collecting samples from the culture effluent, and determining relative fractions using pyrosequencing as before. We found an excellent agreement between experimental data and model simulations (Figure 5), indicating that the fitness benefit of the CcpA-Met19 mutation is growth-rate independent.

**Figure 5.**
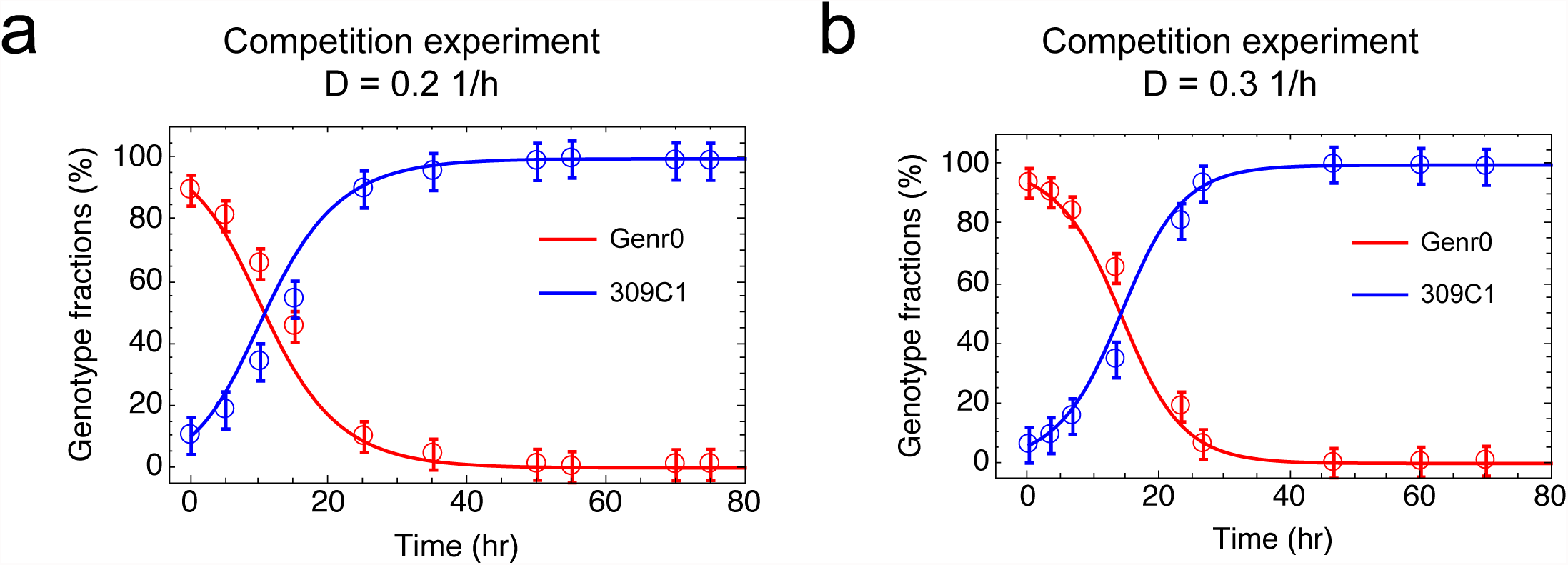
Chemostat competition experiments. Experiments were carried out between Genr0 (red) and 309C1 (blue) at a dilution rate of 0.2 h^-1^ (panel a) and 0.3 h^-1^ (panel b). Marks represent averages and standard deviations based on experimental data. Line depicts results from *in silico* simulations using the model presented here parameterized based on the population dynamics observed during the evolution experiments at a D = 0.5 h^-1^ (Supplementary Figure 13).

## DISCUSSION

### Replicate chemostats show high evolvability and low plasticity

We studied the molecular mechanism of the adaptive strategy employed by *L. lactis* during growth in a glucose-limited chemostat. In this way, we kept the identity of the selective pressure constant, but varied its strength by imposing different dilution rates. Miniature chemostats were constructed and growth medium and protocols developed that allowed highly standardized, parallel and stable conditions for evolutionary studies in quadruplicate for hundreds of consecutive generations, providing true biological replicates of this early evolutionary process.

Intriguingly, we found rapid fixation of exactly the same beneficial amino-acid substitution across all parallel evolutionary experiments, independent of growth rate. This demonstrates a high evolvability with low evolutionary plasticity under these conditions. Evolutionary plasticity is related to the existence of alternative mutations that give rise to a comparable fitness enhancement for a particular selective pressure (Snell-Rood *et al.*, 2010; Forsman, 2014). Plasticity is high when many alternatives exist. Evolvability considers the rate of evolution. It is both related to the number of mutations required, and their fixation rate, to reach a specific quantitative change in fitness. If several mutations are required in sequence for a specific quantitative change in fitness then evolvability is low, and vice-versa. Evolvability can be assessed by determining the number of generations that it takes fitter mutants to fix (Colegrave and Collins, 2008; Pigliucci, 2008). High plasticity and high evolvability occur when alternative mutants fix at similar rates. High evolvability can also occur at low plasticity: then one mutation fixes across replicates and dilution rates. The latter was clearly the case in our study, and it appears that there is a single-peaked fitness landscape, suggesting a strong preference for this specific mechanism, independent of the strength of the selection pressure. As we will discuss shortly, our data suggest that alleviating constraints on membrane occupancy may be that mechanism.

### Mutation in a global regulator rather than specifically in a glucose transporter

In the chemostat, fitter mutants outgrow their competitors because they can grow at a rate equal to the dilution rate, but at a lower concentration of the limiting nutrient (Gresham and Hong, 2014; Bachmann *et al.*, 2016; Wortel *et al.*, 2016). So, in glucose-limited chemostats, one expects enhanced selective pressure toward increased efficiency of glucose uptake, which could be achieved by mutating or elevating the expression of high-affinity transport systems, as previously observed (Bachmann *et al.*, 2012; Solopova *et al.*, 2012). However, although glucose transport activity did increase, we found a mutation with pleiotropic effects in the global carbon metabolism regulator CcpA, rather than in any of the three glucose transporters.

The DNA binding domain of CcpA is highly conserved and in natural isolates of *L. lactis,* as well as in CcpA homologues of other Gram-positive organisms such as *B. subtilis,* the Met residue is present at an equivalent position. Based on the high-resolution structures of CcpA from *B. subtilis*, this residue directly interacts with the nucleobases in the *ackA2* and *glntR* promoter sequences but not with the synthetic *cre* site (Schumacher *et al.*, 2011). Whether CcpA binds with a higher or lower affinity to certain *cre* sites has been investigated in *B. subtilis* (Marciniak *et al.*, 2012). While the focus was on the *cre* site sequence itself, our study gives insights into the effect of the CcpA amino acid sequence on DNA binding. The direct-binding data show that the effect of the amino acid substitution consistently results in a tighter binding of CcpA to *cre* sequences, although not always to the same extent. Other factors such as distance of the *cre* site to the transcriptional start site, and/or the presence of other transcriptional regulators governing the expression of the operon are known to affect gene expression (Deutscher, Francke and Postma, 2006). The different extent of tighter binding of the evolved CcpA, in conjugation with the different regulatory effects that CcpA may adopt for different operons, results in the reprogramming of the this regulon, particularly around membrane proteins (underlying reasons are addressed below). This allows the overall fine-tuning rather than the optimization of a particular trait (e.g. increasing Ks of a single transporter), in response to imposed selection pressure.

CcpA was not the only global regulator to accumulate mutations. We also found a Phe substitution for Ile^214^ in the DNA binding domain of CodY, a global regulator of nitrogen metabolism in *L. lactis*. This mutation was only found in one chemostat but, similar to the mutation accumulated in *ccpA,* indicates a strategy of modulation of global gene expression in response to the culture conditions.

### Fitness enhancement independent from growth rate requires phenotypic plasticity

The fact that we find a single mutation and that we can capture the population dynamics of the CcpA-Met19 allele wash-in at different dilution rates with a simple growth model with fixed kinetics, shows that the mechanism underlying the adaptation to a constant glucose-limiting environment is independent of the growth rate. Yet different growth rates do require different fluxes of glucose uptake, catabolism and biosynthesis. In view of the prevailing perspective of growth-rate dependent regulation of the proteome, including global regulatory mechanisms that may depend on this flux (Kotte, Zaugg and Heinemann, 2010; Basan *et al.*, 2015), this is far from trivial. This seems only possible if the cells possess a form of non-genetic phenotypic plasticity that tolerates the adjustment of the gene-regulatory circuitry under different conditions. A recent functional genomics study of the parent Genr0 strain at different growth rates – using the same standard cultivation conditions – indeed suggests that regulation at the metabolic level is the dominant form of phenotypic plasticity (Goel, Eckhardt, Puri, Jong, Branco dos Santos, Giera, Fusetti, Vos, Kok, Poolman, Molenaar, Kuipers and Teusink, 2015), *i.e.* the proteome of *L. lactis* hardly changed in the wide range of growth rates also studies here (0.2 – 0.6 h^-1^). The presence of such metabolic phenotypic plasticity seems therefore a requirement for our finding of low genetic plasticity and high evolvability. It shows that the intracellular proteome does not impose major constraints on the organism under these conditions, but rather the proteome of the membrane, as we will discuss next.

### Reprogramming of gene expression hints at an important constraint at the level of membrane occupancy

While the mutation in CcpA does increase the expression of genes for high affinity glucose PTS systems (e.g. PTS^Man^), it simultaneously downregulates the expression of unused membrane proteins such as transporters for other sugars and respiratory proteins. This dual role was shown experimentally to indeed result in evolved cells with a more efficient glucose uptake, and suggests that the most successful adaptation to constant glucose-limitation in *L. lactis* requires reprogramming of membrane occupancy, not only upregulation of the specific transporter. The idea that constraints related to limited resources and physicochemical properties affect evolutionary strategies and create trade-offs is much under debate in recent literature (Molenaar *et al.*, 2009; Scott *et al.*, 2014; Goel, Eckhardt, Puri, Jong, Branco dos Santos, Giera, Fusetti, Vos, Kok, Poolman, Molenaar, Kuipers, Teusink, *et al.*, 2015; O’Brien, Utrilla and Palsson, 2016). One interpretation of our data is that the down-regulation of membrane processes provides more membrane space for glucose transporters, in analogy with re-allocation of cytosolic proteins, as happens in *E coli* under different growth conditions (Hui *et al.*, 2015).

Our results are consistent with a detailed computational study of *E. coli* that suggested the presence of different growth regimes with different constraints limiting growth rate (O’Brien *et al.*, 2013). At low growth rates, nutrient uptake was assumed to be the dominant constraint, and cytosolic protein machinery for growth is in excess; only at nutrient excess, internal proteome constraints became dominant. Such excess of cytosolic proteins at sub-maximal growth can explain the predominantly metabolic regulation of flux under our conditions, and hence explains the required phenotypic plasticity (Goel, Eckhardt, Puri, Jong, Branco dos Santos, Giera, Fusetti, Vos, Kok, Poolman, Molenaar, Kuipers and Teusink, 2015). We thus provide an experimental case-study that suggests that indeed uptake constraints are the main constraint for fitness in the chemostat, and that global regulators may be able to overcome them, through pleiotropic effects.

In line with this, we found that the growth rate of the CcpA-Met19 mutant was lower than in wildtype when grown in batch (glucose excess), despite higher glucose uptake capacity - something that is rather common for microorganisms evolved to nutrient-limited conditions (Gresham and Hong, 2014). Also serial batch growth of *L. lactis* in emulsion– a method to select for high cell number – selected for a mutation in the glucose transporter itself, rather than in a regulator thereof (Bachmann *et al.*, 2013). In that case, the mutation reduced glucose transport activity, thereby mimicking a low-glucose environment associated with acetogenic metabolism, a high ATP-yield strategy.

In conclusion, our work shows that evolution to constant growth conditions can be mediated by global regulators such as the carbon catabolite protein CcpA. Metabolic engineering through the mutation of global regulators was first demonstrated by Stephanopoulos and coworkers (Moxley *et al.*, 2009). We here show that the subtle manipulation of global regulators to change entire metabolic networks is a strategy already employed by nature.

## MATERIAL AND METHODS

### Bacterial strains, media and growth conditions

*Lactococcus lactis* subsp. *cremoris* MG1363 was grown in a chemically defined medium developed for prolonged cultivation, CDMPC (Supplementary Table 1). The major differences compared to previously published chemically defined media for *L. lactis* (Thomas, Ellwood and Longyear, 1979; Poolman and Konings, 1988; Jensen and Hammer, 1993) are (i) the implementation of standardized procedures that reduce variations between media batches, (e.g. safeguarding solubility); (ii) the removal of components that are redundant or cause technical (downstream) difficulties or perturbation of cell behavior; and (iii) careful tuning of the concentration of medium compounds. The media consists of a phosphate buffer at pH 6.5, supplemented with all 20 amino acids, the vitamins biotin, DL-6,8-thioctic acid, D-pantothenic acid, nicotinic acid, pyridoxal hydrochloride, pyridoxine hydrochloride and thiamine hydrochloride, and trace metals (NH_4_)_6_Mo_7_O_24_, CaCl_2_, CoSO_4_, CuSO_4_, FeCl_2_, MgCl_2_, MnCl_2_ and ZnSO_4_. Glucose was added to a final concentration of 25 mM and cultivation was performed under an anaerobic headspace. In order to study the effects of long term cultivation under constant conditions, *L. lactis* MG1363 was cultivated in 4 chemostats run in parallel at 30°C at the following dilution rates: 0.5 h^-1^ for a total of 309 consecutive volume changes; 0.6 h^-1^ for over 800 consecutive volume changes. A small sample was removed every 10-15 generations and stored at -80 °C in 20% (v/v) glycerol. In addition, the effluent was collected for optical density at 600 nm (OD_600_) measurements and HPLC analysis.

For subsequent studies, the glycerol stocks were used. 5 ml of fresh CDMPC was inoculated from the frozen glycerol stocks and grown for 16 h at 30°C. The overnight culture was subsequently diluted to a starting OD_600_ of 0.025-0.050 in either CDMPC or M17 medium (Oxoid, Basingstoke, United Kingdom) supplemented with 0.5% (w/v) glucose or other carbon sources as indicated.

### Chemostat competition experiments between Genr0 and 309C1

The growth model parameterized using the data from the evolution experiments at a D of 0.5h-1 was used to designed the experiments, such that the initial fraction and sampling time-scheme would be adequate to capture the predicted wash-in kinetics of the evolved cells. Actual competition experiments were preceded by running four-parallel chemostats for 10 volume changes as described above for Ds of 0.2 and 0.3h-1. Of the four steady states independently established for each D, three included Genr0 and one 309C1. The competition stage was then initiated by transferring the 309C1 culture into the Genr0 steady states such that they made up ~10% of the population. After proper mixing a population sample is taken and considered to be t0. Subsequently, and according to a predetermined sampling scheme, samples are removed from the chemostat effluent. The relative fractions of the population corresponding to Genr0 and 309C1 were then determined in time resorting to pyrosequencing the *ccpA* locus (detailed in the supplementary material).

### Whole-genome sequencing and mutation detection

DNA was extracted from the *L. lactis* single isolates using the GenElute™ Bacterial Genomic DNA kit (Sigma-Aldrich, St. Louis, MO, USA) according to the manufacturer’s instructions except that the cells were first pre-treated with lysozyme at 10 mg/ml for 1 hour at 50°C. The isolates were sequenced with 100bp single-end reads across one lane of an Illumina HiSEQ 2000 flow cell (Illumina). The resulting reads were deposited in the NCBI-SRA database (PRJNA185994). Reads were mapped to the reference genome of *L. lactis* ssp. *cremoris* MG1363 (GenBank accession number AM406671) and mutations detected using the CLC BIO Genomic Work Bench Suit 4.5 (CLC BIO, Arhus, Denmark). The mutation lists were verified using PCR amplification and subsequent Sanger sequencing.

### Microarray analysis

For RNA extraction, all strains were grown in CDMPC to the midexponential growth phase. Total RNA was isolated using the High Pure RNA isolation Kit (Roche Applied Science) and the quality was tested using an Agilent Bioanalyzer 2100 (Agilent Technologies Netherlands BV). All procedure regarding microarrays were performed as described previously (van Hijum *et al.*, 2003, 2005). Data was processed and normalized using the Lowess method with Micro-Prep (van Hijum *et al.*, 2003) and lists of significant gene expression changes were generated on the basis of a Bayes *p-*vale and Benjamini Hochberg multiple testing corrections (Hochberg and Benjamini, 1990). The DNA microarray data have been submitted to GEO with accession number GSE67657.

The expression data was further analyzed by the Rank Product analysis (Breitling *et al.*, 2004) using the software package R version 2.10.1. This generates a list of genes ranked according to ln ratio and calculates a conservative estimate of the percentage of false positives (pfp). Proteins with pfp values smaller than 0.05 (5%) were regarded as differentially expressed. The lists of significant gene expression levels were subjected to functional class analyses supported by the Genome2D webserver (http://server.molgenrug.nl). This results in a list of classes that are overrepresented in the dataset supplied on the basis of a hypergeometrical distribution test. The data for the functional classes was derived from the KEGG public data base (http://www.genome.jp/kegg/ko.html) and for regulons from PePPER (de Jong *et al.*, 2012).

### [^14^C]-glucose uptake

Glucose uptake was monitored in whole cells using [^14^C]-labeled glucose essentially as previously described (Gouridis *et al.*, 2015). Briefly, all strains were grown in CDMPC and harvested at the mid-exponential growth phase, washed twice in ice-cold KPi buffer (50 mM, pH 6.5) and resuspended in KPi buffer to an OD_600_ of 0.5. Uptake assays were performed at 30°C with stirring. [^14^C]-glucose was added to 100 μl cell suspension aliquots to a final concentration of 1.2-200 μM (0.005 μCi). The reactions were stopped by the addition of 2 ml of ice-cold 0.1 M LiCl and the samples were collected on 0.45μm pore-size cellulose nitrate filters (Schleicher and Schuell GmbH, Dassel, Germany). The filters were again washed with 2 ml 0.1 M LiCl. The background was determined by adding the radiolabeled substrate to the cell suspension immediately after the addition of 2 ml of ice-cold LiCl, followed by filtering. The assays were performed in triplicate using cells from two independent cultures.

### Electrophoretic mobility shift assays

To determine the *cre* site DNA binding affinity of CcpA, binding assays were performed with Cy3-labeled oligonucleotides. DNA oligonucleotides were from Biolegio B.V. (Nijmegen, the Netherlands) with the (+) strand labelled at the 5’end with Cy3 and the (-) strand unlabeled. The oligonucleotides were mixed 1:1, incubated at 98°C for 3 minutes and allowed to cool slowly to room temperature to facilitate annealing. The annealed DNA was used at 4 nM and added to binding buffer (20 mM Tris-HCl pH8, 100 mM KCl, 10 mM MgCl2, 1 mM EDTA, 1 mM DTT, 5% glycerol) with additional BSA to 40 μg/ml. CcpA was added to concentrations between 100 nM and 800 nM. The binding reaction was performed at 30 °C for 20 minutes after which the samples were mixed with 5 X gel loading buffer (0.25% bromophenol blue, 40% sucrose). The samples were analysed by electrophoresis on 7.5% native polyacrylamide gels using a TBE buffer (90 mM Tris, 90 mM Boric Acid, 2 mM EDTA pH8.0). The fluorescent-labelled oligonucleotides were visualised on a LAS-4000 imager (Fujifilm).

### Molecular dynamics simulations

The simulated structures were based on the crystal structures of CcpA bound to different DNA sequences by Schumacher *et al*. (Schumacher *et al.*, 2011). Homology models of *L. lactis* proteins were created using the Swiss Model Server (Arnold *et al.*, 2006). The sequence identity of the DNA binding domain in proteins from *B. subtilis* and *L. lactis* is 77 % and only this region of the protein was used for the simulation. Structures with differing DNA sequences were produced by fitting the differing bases to a crystal structure before equilibrating the system. The protein domains after residue 64 were removed from the simulation system and a harmonic bond with a 1000 kJ/(mol nm^2^) force constant was set between the backbone atoms of the last residues to restrain the structure of the remaining domains. Each system was solvated in a dodecahedron box where each face of the box was at least 1 nm from the protein and the DNA. This amounted in average to about 14500 water molecules, 130 Na^+^ ions and 110 Cl^-^ ions, which corresponds to a 420 mM NaCl solution and counter-ions.

All simulations were run using the Gromacs 4.5.5 (Hess *et al.*, 2008) simulation package. The protein and DNA were modelled with CHARMM27 force field with CMAP corrections (Foloppe and MacKerell Jr., 2000; MacKerell, Feig and Brooks, 2004) together with the original TIP3P water model (Jorgensen *et al.*, 1983) as implemented in Gromacs 4.5.5.

Free energy changes were calculated from simulations using the Bennett acceptance ratio method (Bennett, 1976) with the g_bar tool in Gromacs. Gromacs is a free software package available at http://www.gromacs.org/. Relative free energies of binding were calculated using a thermodynamic cycle described in Supplementary Figure 9.

## ACKNOWLEDGEMENTS

This study was funded by the Kluyver Centre for Genomics of Industrial Fermentation and the Netherlands Consortium for Systems Biology (NCSB), within the framework of the Netherlands Genomics Initiative (NGI)/NWO. FBS, FJB and BT are supported by the Netherlands Organization for Scientific Research (NWO) through VENI grant 863.11.019, VIDI grant 864.11.011 and VICI grant 865.14.005 respectively.

## Supplementary Results

### Mutations in all sequenced strains compared to published sequences

The sequences of Genr0 and the evolved strains were compared to those published for *L. lactis* MG1363. All strains contained a mutation in *malR* which would results in a frameshift in the maltose operon transcriptional repressor MalR. This could be due to sequencing errors from adjacent C nucleotides or a truncated MalR protein. The truncation would result in a HTH DNA binding domain without a ligand-binding domain.

### Mutations unique to strains 309C2, 309C3 and 309C4

The effect of the mutations identified in *rpoC* in 309C2 and *greA* in 309C4 on transcriptional fidelity were investigated using *lacZ* constructs with a premature stop codon at position Glu^13^ (P. Gamba and J.W. Veening, unpublished data). However, β-galactosidase activity was similar in the evolved strains and the original strains Genr0 and *L. lactis* MG1363 (Supplementary Figure 2), indicating that in the evolved strains, RNA polymerase and elongation factor function as they do in wild type strains.

Since proteins encoded by llmg_1597, a hypothetical protein possibly with RNA-binding function, and *llrF,* a two-component system regulator, are not well-characterized, no prediction as to change or function could be made for the mutations identified. In contrast, CodY is a well-studied transcriptional regulator of nitrogen metabolism and based on structural studies it can be predicted that the SNP identified results in a Phe substitution for Ile^214^ in the DNA binding domain (Levdikov, Blagova, Joseph, Sonenshein, & Wilkinson, 2006). The absence of a high-resolution structure of CodY bound to a DNA operator makes any further prediction of the effect on DNA binding difficult.

A number of SNPs were predicted to lie in non-coding regions at positions that lie in or near putative promoter sequences, thus making predictions as to the effect of such mutations difficult. The SNP at position 490263 resulted in an altered ribosomal binding site (GGAGGA to GGAGAA) upstream of the gene for a Hu-like DNA binding protein, *hllA*. This could suggest that a modest reduction of the translation of this mRNA, encoding a putative DNA-binding protein, could take place.

### Global gene expression patterns of evolved strains

The evolved strains were revived in CDMPC and grown until mid-exponential phase at which microarray analysis using in-house *L. lactis* MG1363 slides (Kuipers et al. 2002) was performed. Gene expression in the evolved strains was compared to that of Gen0. A total number of 377 genes exhibited changed expression in the evolved strains (Supplementary Table 4). Approximately a third of the changes were in at least 3 of the 4 strains. Altered gene expression was dominantly in genes involved in membrane transport, especially carbohydrate import (Supplementary Figure 3A, Supplementary Table 3, Supplementary Table 4). Regulon analysis of the differentially expressed genes revealed that CcpA-regulated genes were over-represented in the data set. For strain 309C4, those regulated by CodY were prevalent as well (Supplementary Figure 3B, Supplementary Table 3).

CcpA controls the preferential use of glucose over other sugars (Deutscher et al. 2006), including expression of sugar uptake systems such as those for glucose. Transcriptome analysis of all the evolve strains revealed similar patterns of expression and increased glucose uptake (Supplementary Figure 3C and D).

The ability to utilize other carbon sources was diminished for the evolved strains (Supplementary Figure 4A) in accordance with the down regulation of, amongst others, genes encoding for maltose transporters, ABC sugar import systems and sugar utilization enzymes in the microarray data (Supplementary Table 3). Many of these genes are CcpA-regulated.

### Phenotypic characterization of evolved strains to identify effects of transcriptional regulation

The strains were not only adapted to the constant culture conditions through expression changes in carbohydrate uptake and utilization systems, but also through changes in the expression levels of genes involved in amino acid uptake and metabolism. Ordinarily, *L. lactis* MG1363 grows faster when amino acids are taken up as oligopeptides than when only free amino acids are available (Supplementary Figure 5B). This is due to the efficient import of oligopeptides by the major oligopeptide transporter Opp (Detmers et al., 2000). Cultivation in free amino acids led to the down regulation of the entire *opp* operon (Supplementary Figure 5A) and the cells were as a result impaired in growth on oligopeptide containing medium (Supplementary Figure 5B and C). The difference in growth in oligopeptide-containing medium between the evolved strains and Genr0 is well correlated with the expression levels of the *opp* operon except in the case of 309C4. In this strain the down regulation of the *opp* operon appears to be partially compensated for by the concomitant up regulation of the analogous *opp2* operon. This operon was previously thought to be cryptic in *L. lactis* MG1363 (Sanz et al. 2004), but in our evolved strain is expressed and OppA2 was detected using mass spectrometry (data not shown).

Prolonged cultivation under an anaerobic atmosphere had a dramatic effect on the ability of the evolved strains to utilize the electron transport chain present in *L. lactis*. *L. lactis* is unable to synthesize heme *de novo,* but if heme is added to the growth medium, the bacterium can synthesize the cytochromes needed for the generation of a proton motive force (PMF) and subsequent ATP synthesis at the membrane-embedded ATP synthase (Brooijmans et al. 2007). This results in an increase in biomass (Cesselin et al. 2001) as observed for Genr0 (Supplementary Figure 6). The evolved strains did not show any increase in biomass upon addition of heme to the medium and expression data showed the down regulation of a number of NADH oxidases/dehydrogenases (Supplementary Table 3) which are likely to be responsible for the inability of these strains to respire.

In addition the evolved strains displayed a well-characterized long-chain phenotype associated with a deficiency in the major autolysin AcmA (Supplementary Figure 7) (Buist et al. 1995). The expression of the *acmA* gene was down regulated in all the evolved strains (Supplementary Table 3) and in agreement with the down regulation of the *acmA* gene, the evolved cells were less prone to lysis in the stationary growth phase (Supplementary Figure 7).

### Effect of M19I mutation on the binding affinity of CcpA

The effects of the *ccpA* mutations were investigated using homology modeling, molecular dynamic simulations and *in vitro* binding assays using purified CcpA from the mutated strains. The high-resolution structure of *B. subtilis* CcpA was used to model *L. lactis* CcpA (Figure 4A and Supplementary Figure 8) (Schumacher et al. 2011) since the one published for *L. lactis* CcpA lacks the N-terminal DNA-binding domain (Loll et al. 2007). The structure from *B. subtilis* includes CcpA bound to the DNA catabolic response element (*cre* site) as well as the activating phosphoprotein HPr. Met19 is located in the second helix contained in the DNA binding domain. The methionine at this position is highly conserved across all Gram-positive CcpA proteins sequenced to date and in 4 natural isolates of *L. lactis* for which the *ccpA* locus was resequenced (Supplementary Table 6).

The binding of CcpA from Genr0 and 309F1 to 4 DNA operators was examined: synthetic *cre* sites, *cre* site upstream of *ptcB*, *cre* site upstream of *mtlD* and, as a negative control, the CodY binding sequence upstream of *oppD* (Figure 4D and Supplementary 10), representing genes that are down regulated as well as up regulated in the evolved strains. For all the cre sites tested, the CcpA-Ile19 variant showed an increase in binding affinity for the DNA operators tested. This was most pronounced for the sequence upstream of *ptcB* and least pronounced for *mtlD*.

### Evolution of Ccp-Thr^19^ variant

The appearance of and subsequent dominance of the *ccpA* mutations was tracked in time during the evolution experiment through PCR amplification of the *ccpA* locus from the frozen stocks. The PCR products were resequenced using traditional Sanger sequencing, with manual chromatogram inspection, as well as pyrosequencing, which allows for the quantitation of SNPs in a population. In addition, frozen stocks were streaked onto CDMPC agar plates and the *ccpA* locus was PCR amplified and resequenced from single colonies. The CcpA-Ile^19^ variant emerged around 50 generations in the evolved strains and by generation 150 was the dominant population in the chemostat (Supplementary Figure 11 and 12). The pattern of emergence was different for strain 309C4, with the emergence of a second variant, i.e. CcpA-Thr^19^. This variant was only detected after 250 generations and by 309 generations both CcpA variants existed in the population. Sequencing of numerous single colony isolates could not detect any wild type *ccpA* by 300 generations (data not shown).

Sequencing of single colony isolates from the C4 frozen stocks, revealed that the unique CcpA-Thr^19^ variant is linked to the CodY-Phe^214^. At 309 generations the only combinations identified were CcpA-Ile^19^ and CodY-Ile^214^ (i.e. wild type CodY) and CcpA-Thr^19^ and CodY-Phe^214^. Stocks resequenced from earlier in the evolution experiment also contained CcpA-Thr^19^, CodY-Ile^214^ pairs, indicating that the *codY* mutations occurred in the unique CcpA-Thr^19^ background.

### Model description and parameter estimation

The differential equations describing the rates of the change in the limiting nutrient concentration (s) and the biomass concentrations of the wild type (w) and the mutant (m) are,

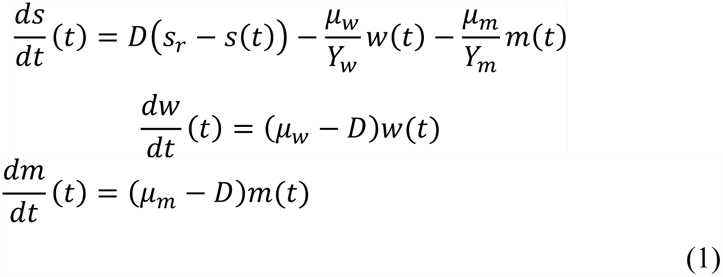

with: 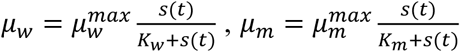, D as the dilution rate in h^-1^, μ as the specific growth rate in h^-1^, Y is the biomass yield in gDW.mM^-1^, μ/Y as the specific nutrient uptake rate in mM.gDW^-1^.h^-1^, K as the Monod constant in mM, s_r_ as the vessel concentration of the limiting nutrient in mM, s as the reactor concentration of the limiting nutrient in mM, and w and m as the biomass abundances in gDW.L^-1^. The subscripts “w” and “m” indicate whether the parameter concerns the wild type or the mutant. Alternatively, we could have taken the OD to represent the biomass concentration, which we now take as gDW.L^-1^. Considering the experimental data it is more convenient to consider a dimensional time defined as *τ* = *Dt*, which is what we call generations. With this definition we obtain,

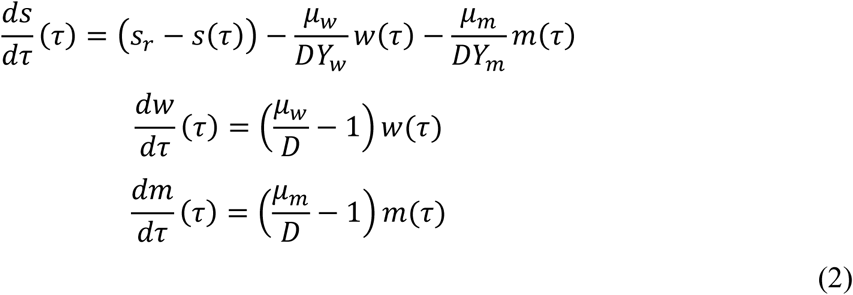

Since we are considering the fixation of the mutant, the initial conditions are given by the steady state of wild type in the absence of the mutant:

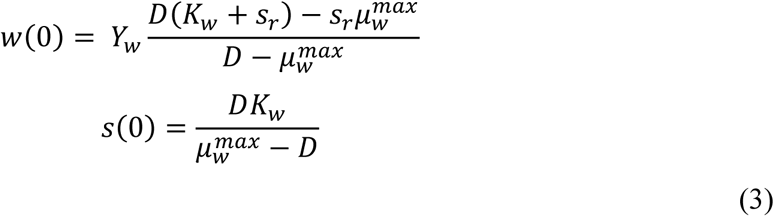

Equations 2 and 3 were fitted against the experimental data. The results of the fit are shown in Supplementary Table 1 and Supplementary Figure 13. These data indicate that observed experimental data is in agreement with a basic chemostat model of the competitive growth of two microorganisms. The values of the fitted parameters are not the only set of values that describe the data. The data set is too small, and hence more variables would have to be measured to obtain a unique estimation of parameter values.

## Supplementary materials and methods

### Chemically defined medium for prolonged cultivations

*Lactococcus lactis* subsp. *cremoris* MG1363 was grown in a chemically defined medium developed for prolonged cultivation, CDMPC (Supplementary Table 1). The media consists of a phosphate buffer at pH 6.5, supplemented with all 20 amino acids, the vitamins biotin, DL-6,8-thioctic acid, D-pantothenic acid, nicotinic acid, pyridoxal hydrochloride, pyridoxine hydrochloride and thiamine hydrochloride, and trace metals (NH_4_)_6_Mo_7_O_24_, CaCl_2_, CoSO_4_, CuSO_4_, FeCl_2_, MgCl_2_, MnCl_2_ and ZnSO_4_. The composition was based on the nutrient requirements and biomass composition of *L. lactis* MG1363. The major differences compared to previously published chemically defined media for *L. lactis* (Thomas et al. 1979; Poolman and Konings 1988; Jensen and Hammer 1993) are (i) the removal of acetate and ammonium ions; (ii) removal of redundant and/or unessential vitamins such as vitamin B12, or those that could in addition impose (downstream) technical challenges such as riboflavin, which fluoresces, and folate, which is difficult to dissolve and likely to precipitate during media preparation; (iii) removal of trace elements that are likely to precipitate; (iv) removal of all non-essential nucleic acid precursors; (v) adjustment of phosphate concentration such that it provides enough buffering capacity to enable growth until OD_600_ of 0.8, while not inhibiting growth rate; (vi) adjustment of amino acid composition to 2.5-fold the requirement for 1gDW·L^-1^ according to the biomass composition (Oliveira et al. 2005); and (vii) implementation of a new protocol that reduces variations between media batches by avoiding precipitation and heat or light degradation. Glucose was added to a final concentration of 25 mM and cultivation was performed under an anaerobic headspace.

### Standardized cryopreservation and inoculation procedure

The outcome of laboratory evolution experiments can be greatly affected by the composition of the population used to seed each of the prolonged cultivations. In order to minimize background genetic variation and variations in pre-culture condition, we developed a standardized cryopreservation and inoculation procedure inspired by industrial procedures to create, maintain and distribute single strain isolate stocks (Figure 1, main text). All the prolonged cultivations in this study were carried out from the working stocks generated via this procedure.

#### Single colony isolation and parent-stock preparation

A glycerol stock of *L. lactis* MG1363 (provided by Jan Kok, University of Groningen, the Netherlands) was used to inoculate M17 enriched with 0.5% glucose (GM17). This culture was streaked out from GM17 onto CDMPC agar plates (CDMPCA) and incubated for 48 hours at 30° C. An isolated colony was transferred to CDMPC and incubated for 24 hours at 30° C. A 1% inoculum of the freshly grown culture was used to inoculate CDMPC. After 5 consecutive transfers under the same conditions and using 1% inoculum to seed subsequent cultures, the resulting culture was streaked out in CDMPCA and incubated for 24 hours at 30° C. An isolated colony was transferred to 7.5 ml of CDMPC and incubated for 24 hours at 30° C reaching a final OD_600_ of 1.165. “Parent-stocks” were then prepared by adding 3.75 ml of sterile 60% glycerol and dividing the mixture in 11 aliquots of 1 ml in self-standing Nunc™ Cryotubes (366656), kept at -80° C for at least for 48 hours before reviving or moving.

#### Working-stock preparation

One of the 11 aliquots of the parent-stock was used to inoculate 199 ml of CDMPC (0.5% inoculum) and incubated for 20 hours at 30° C, until the culture was full-grown. 100 ml of sterile 60% glycerol was then added, resulting in 300 ml of 20% glycerol stock of *L. lactis* G0 in CDMPC. “Working-stocks” were then prepared by dividing the latter in 960 aliquots of 0.3 ml in 1.2 ml sterile polypropylene cluster tubes 96-well racked from Corning Incorporated Costar^®^ (4413). These were capped using 8-Cap Strips from Corning Incorporated Costar^®^ (4418) and separated into individual vials using a slightly heated bistoury and ruler. Working-stocks were stored at − 80° C for at least 24 hours before reviving or moving.

#### Standardized inoculation procedure

A working-stock was revived by thawing a 0.3 ml aliquot of 20% glycerol stock of *L. lactis* G0 in CDMPC and used to inoculate 44.7 ml of CDMPC (0.67% inoculum). This culture was incubated for 16 hours at 30° C, which corresponds to the time at which the culture reaches early stationary phase (OD_600_ ~1.00±0.15). This freshly-grown culture cultivated in a standardized fashion and seeded from a controlled inoculum, was then used directly as the inoculum of downstream cultivations and assays (chemostats, batch cultures, etc.) performed by different experimentalists across several laboratories committed to working in a standardized way.

#### Moving and storage of cryopreserved stocks

The 11 aliquots of 20% glycerol stock of *L. lactis* G0 in CDMPC (parent-stock) were divided amongst the laboratories of the University of Amsterdam, VU University Amsterdam, Groningen University and Research Center and Wageningen University, preserved in different −80° C freezers within these institutes. The 960 aliquots of 20% glycerol stock of *L. lactis* G0 in CDMPC (working-stock) were divided amongst all the collaborators involved in standardized systems biology studies using *L. lactis* G0 as the model organism. Both the parent- and working-stocks were kept frozen at all times and transported accordingly.

### Experimental set up

*L. lactis* MG1363 was cultivated in 4 chemostats run in parallel at 30°C at the following dilution rates: 0.5 h^-1^ for a total of 309 consecutive generations; 0.6 h^-1^ for a total of 700 consecutive generations. Our design makes use of bioreactor lids that can fit any vessel with a GL45 thread. It allows for two connectors to be inserted at variable depths, used here to control the working volume and sampling. These custom made bioreactors were set-up to make four parallel chemostat cultivations controlling stirring, temperature, pH, flow, volume and gas composition. The newly developed caps are versatile and can be readily adapted to other types of controlled cultivation, such as fed-batch, pH-auxostats, turbidostats, retentostats, amongst others. Culture samples can be withdrawn directly through a port present in the centre of the cultivation vessel or by rerouting the effluent using acetal quick-disconnect couplings (Masterflex via Cole Parmer, Schiedam, Netherlands) Throughout the prolonged cultivations samples were collected from the effluent to leave the cultivation undisturbed.

For each set of parallel chemostats, temperature was controlled using a water bath (Thermo Haake DC30-W13/B via Cole Parmer, Schiedam, Netherlands) set to 30°C. Even stirring was ensured using an immersible stirrer unit connected to a remote controller (Cole Parmer, Schiedam, Netherlands) set to ~300 rpm. Mixing was tested by pulsing with glucose to a final concentration of 60 mM and taking samples at three different positions of the bioreactor. Minor differences were observed between samples harvested at different time points (<5%), and at small-time scales (<10 seconds) the solution appeared equilibrated. pH was measured separately in each culture using a 120 mm gel-filled pH-sensor (Applikon, Schiedam, Netherlands) connected to an Alpha-pH800 (Eutech, Thermoscientific) that controlled a 1 rpm L/S pump drive equipped with an Easy-Load II pump head (Masterflex via Cole Parmer, Schiedam, Netherlands) that titrated the culture with a solution of 2.5 M NaOH. The culture was continuosly flushed through the headspace with a mixture of 5% CO2 and 95% N2 at a flow rate of 10 culture volumes per hour (i.e. 600 ml/h). The dilution rates of the different prolonged cultivations carried out in this study were set by controlling the flow of CDMPC using a variable speed L/S pump drive equipped with four Easy-Load II pump heads (Masterflex via Cole Parmer, Schiedam, Netherlands). All parallel cultivations were supplied with CDMPC from the same medium vessel.

A small sample was removed every 10-15 generations and stored at -80 °C in 20% (v/v) glycerol. In addition, the effluent was collected for optical density at 600 nm (OD600) measurements and HPLC analysis. For subsequent studies, the glycerol stocks were used. 5 ml of fresh CDMPC was inoculated from the frozen glycerol stocks and grown for 16 h at 30°C. The overnight culture was subsequently diluted to a starting OD_600_ of 0.025-0.050 in either CDMPC or M17 medium (Oxoid, Basingstoke, United Kingdom) supplemented with 0.5% (w/v) glucose or other carbon sources as indicated.

### Pyrosequencing

Frozen glycerol stocks served as template material for PCR reactions. The PyroMark platform from QIAGEN was used for pyrosequencing and reactions were performed as per manufacturer’s instructions. The region surrounding the point mutations was amplified and biotinylated with the use of a biotinylated reverse primer. To detect the ratio of SNPs in the population, single stranded primers were designed so as to terminate before the codon containing the mutation.

### Recombinant DNA methods

For cloning purposes, *Escherichia coli* XL1 blue was used. *ccpA* from the evolved *L. lactis* strains was cloned into the pQE30 plasmid essentially as described to create pQE30ccpA (Kowalczyk and Bardowski 2003). This allowed for the placement of 6 X His at the 5’-end of the *ccpA* gene. The *ccpA* gene was PCR amplified from genomic DNA from the single colony isolates from strains Genr0, 309C1, 309C2 and 309C4. *E. coli* XL1blue harbouring pQE30ccpA was used for overproduction of CcpA by IPTG induction. CcpA was purified using Ni-NTA resin (Qiagen, Germantown, MD, USA) as previously described (Kowalczyk and Bardowski 2003). The protein concentration was determined with the DC Protein assay kit (Bio-Rad, Hercules, CA, USA) using BSA as a standard.

### Protein identification by liquid chromatography-mass spectrometry

The samples were separated by SDS-PAGE and stained with Bio-Safe Coomassie (Bio-Rad, Hercules, CA, USA). In-gel tryptic digestion was followed by peptide extraction and LC-MS/MS as described in detail in Drop *et al.,* 2011. The MS raw data was submitted to Mascot (Version 2.1, Matrix Science, London, UK) and searched against the *L. lactis* MG1363 proteome. Protein identifications were based on at least 2 unique peptides identified by MS/MS, each with a confidence of identification probability of at least 95%.

### Molecular dynamics simulations

The free energy of mutating the protein was calculated while protein was bound to DNA and free in solvent and the relative binding free energy of the mutated protein compared to the wild type was obtained by subtracting the free energy of the former from the latter (Supplementary Figure 9).

The free energy of mutating the protein was calculated using a topology with two side chains at residue 19. The two side chains did not interact with each other and the mutation was done in three steps. In the first step, the charges of the wild type methionine side chain were uncoupled from rest of the system while the mutated side chain was completely non-interacting. In the second step, the Lennard-Jones interactions of methionine side chain were uncoupled while at the same time the Lennard-Jones interactions of the mutated side chain were coupled to the side chain. In the third step, the charges of the mutated side chain were coupled to the system while the methionine side chain was completely non-interacting. The first and third steps were performed using 5 simulations with different lambda values while the second step used 11 lambda values. Soft-core interactions were used in the second step with soft-core parameter **α** set to 0.5. An error estimate is calculated from block averaging each 1 ns of the simulations and using the standard deviation of the obtained values as the error.

In the simulations, the protein monomers can move relatively freely in solution when DNA is not present and sampling of conformations becomes an issue. To ensure that the conformations sampled would be similar for all CcpA variants, rotational and translational constraints were used to prevent the monomers from moving with reference to each other as described in (50).

A 2 fs time step was used with all bonds constrained using the LINCS algorithm (Hess et al. 1997). Simulations were run with periodic boundary conditions in all directions. Non-bonded interactions were treated as recommended in (Bjelkmar et al. 2010). Electrostatics were calculated using PME (Essmann et al. 1995) with a 1.2 nm cut-off. Lennard-Jones interactions were switched off between 1.0 and 1.2 nm and dispersion correction was applied to both energy and pressure. The protein-DNA system and the solvent were separately coupled to a 298 K heat bath using the velocity rescaling thermostat (Bussi et al. 2007) with τ_T_ = 0.5 ps. A 1 bar pressure was kept using an isotropic Parrinello-Rahman (Parrinello and Rahman 1981) pressure coupling with τ_p_ = 4.0 ps and 4.5e-5 bar^-1^ compressibility. Each system was energy minimized and briefly equilibrated with position restraints before being equilibrated for 10 ns. After equilibration each simulation was run for 20 ns except the systems with deviant *cre* sites that were simulated for 10 ns.

All simulations were run using the Gromacs 4.5.5 (Hess et al. 2008) simulation package. The protein and DNA were modelled with CHARMM27 force field with CMAP corrections (Foloppe and MacKerell Jr. 2000; MacKerell et al. 2004) together with the original TIP3P water model (Jorgensen et al. 1983) as implemented in Gromacs 4.5.5.

Free energy changes were calculated from simulations using the Bennett acceptance ratio method (Bennett 1976) with the g_bar tool in Gromacs. Gromacs is a free software package available at http://www.gromacs.org/.

### Enzyme reactions

L-lactate dehydrogenase (L-LDH) was purchased from Sigma-Aldrich (St. Louis, MO, USA). L-LDH was added to culture supernatant to a final concentration of 0.2U/ml, NADH was added to 2 mM and the reaction incubated for 15 minutes at 25°C. X-prolyl dipeptidyl aminopeptidase (PepX) activity was measured using the chromogenic substrate Ala-Pro-*p*-nitroanilid (Bachem Feinchemicalien AG, Bubendorf, Switzerland) as described (Buist et al. 1998).

β-galactosidase assays were performed as previously described (Israelsen et al. 1995; Kloosterman et al. 2006).

### Miscellaneous

Oligopeptides were from JPT Peptide Technologies GmbH (Berlin, Germany).

## Supplementary Figures Legends

**Supplementary Figure 1.**
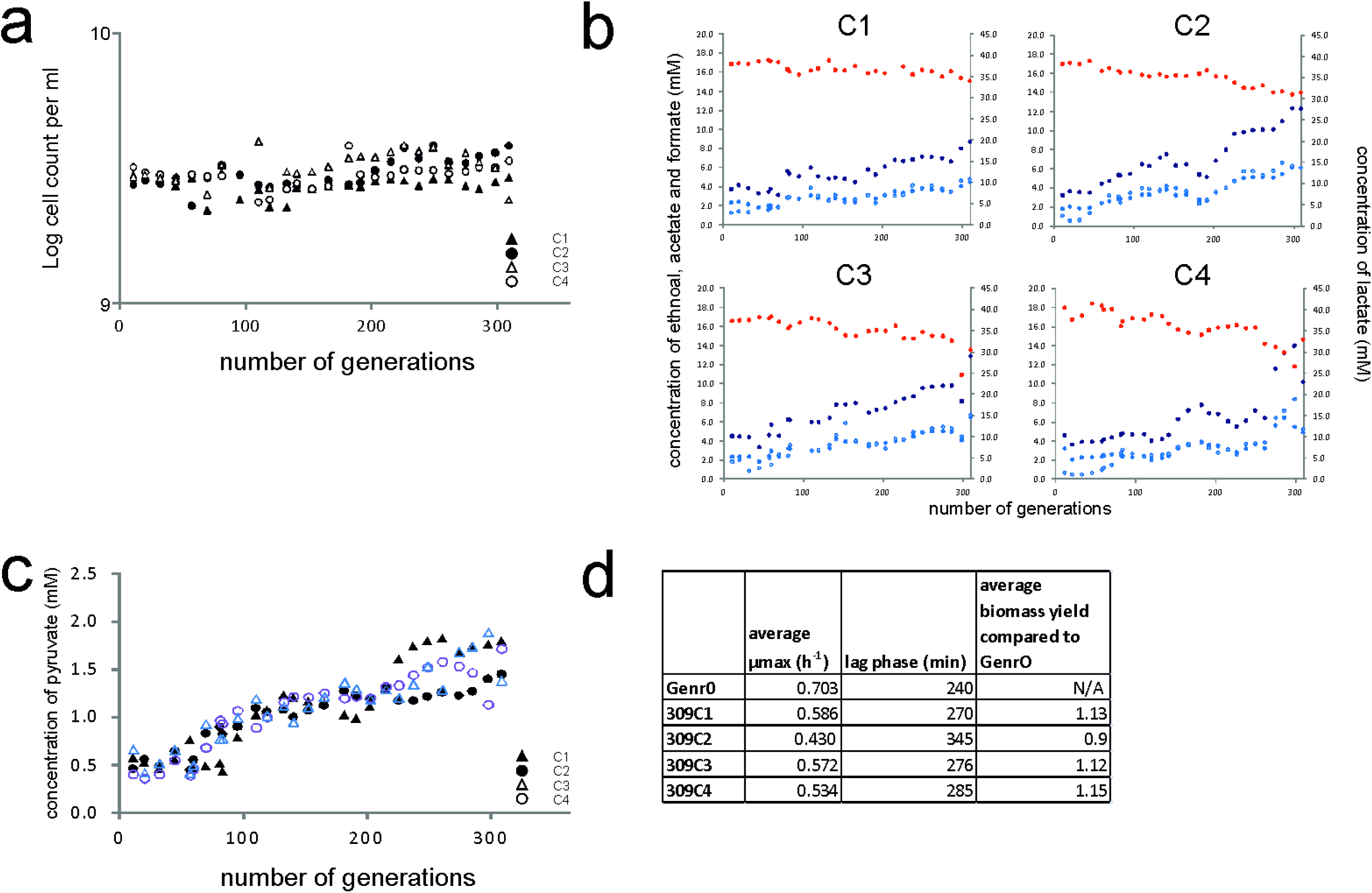
Strain characteristics during the evolution experiments at D of 0.5h-1 and of the resulting evolved strains. For each of the chemostats (labeled C1 to C4) the cell counts (A) and organic acids (B) were measured. The values for lactate (red), formate (dark blue), acetone (blue) and ethanol (light blue) are shown. The levels of pyruvate increased in all the chemostats during the evolution experiment (C). The evolved strains were revived in batch culture and the growth characteristics determined (D).

**Supplementary Figure 2.**
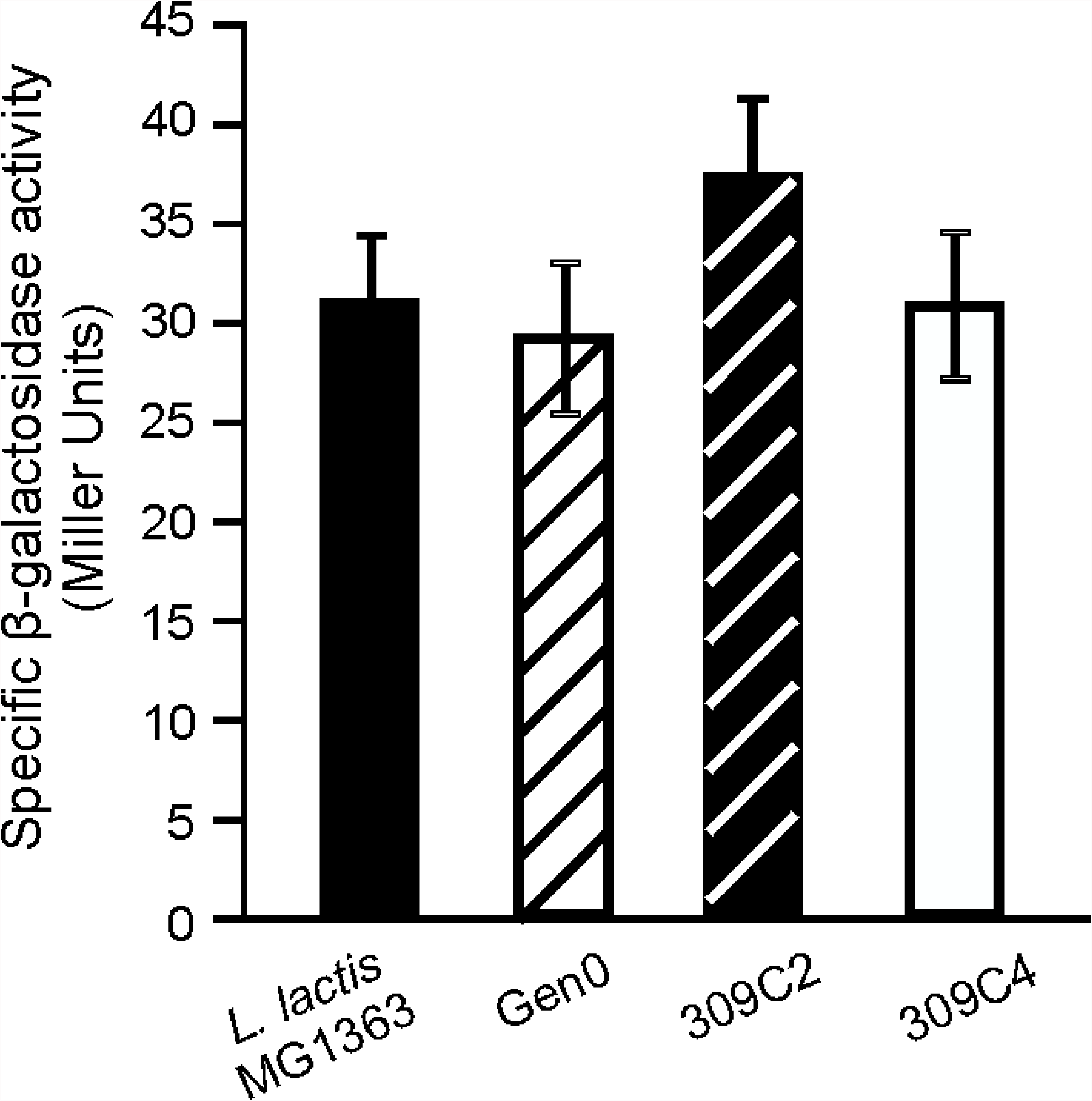
Transcriptional fidelity in evolved strains is comparable to the original ones. Strains were grown in CDMPC and harvested at the mid-exponential phase of growth. Data are averages of three measurements with the error bars indicating the standard deviation. Specific β-galactosidase activity was measured in the strains indicated using pPG6 (P. Gamba and J.W. Veening, unpublished data), which contains a premature stop codon at Glu13.

**Supplementary Figure 3.**
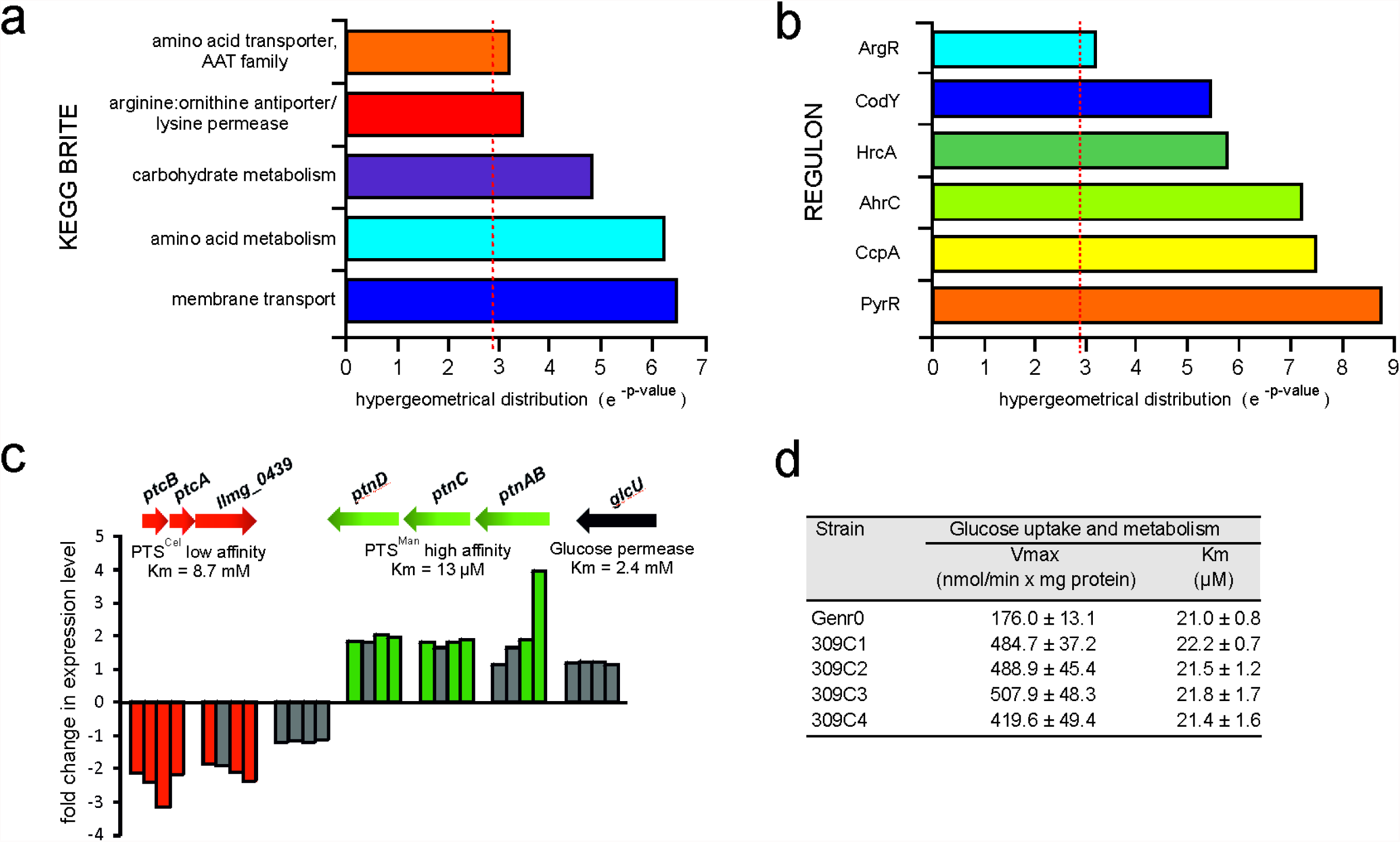
Altered gene expression in evolved strains is focused on transport pathways. Genes were considered to be significantly changes with a Bayes p-value score of less than 0.05 and a pfp value of less than 0.05. Genes found to change significantly in the evolved strains were grouped into functional classes. The p-value is the summation of the hypergeometrical distribution and p-values less than 0.05 were considered to be significant. (a) Over-represented KEGG BRITE classes found. (b) Over-represented regulons found. (c) The expression of genes involved in glucose uptake in *L. lactis* MG1363 were significantly changed in the evolved strains. Highlighted in green are genes that were up-regulated and in red for down-regulated. Gene expression changes not meeting the significance cut-off values are highlighted in grey. (d) Kinetic parameters of glucose transport in Genr0 and the evolved stains were determined in cells grown to exponential phase in CDMPC. Glucose transport was assayed with the use of [14-C]-labeled glucose. Values of three independent experiments were averaged and are reported ±SD. Vmax and Km were determined using glucose concentrations from 1.2 to 200 μM.

**Supplementary Figure 4.**
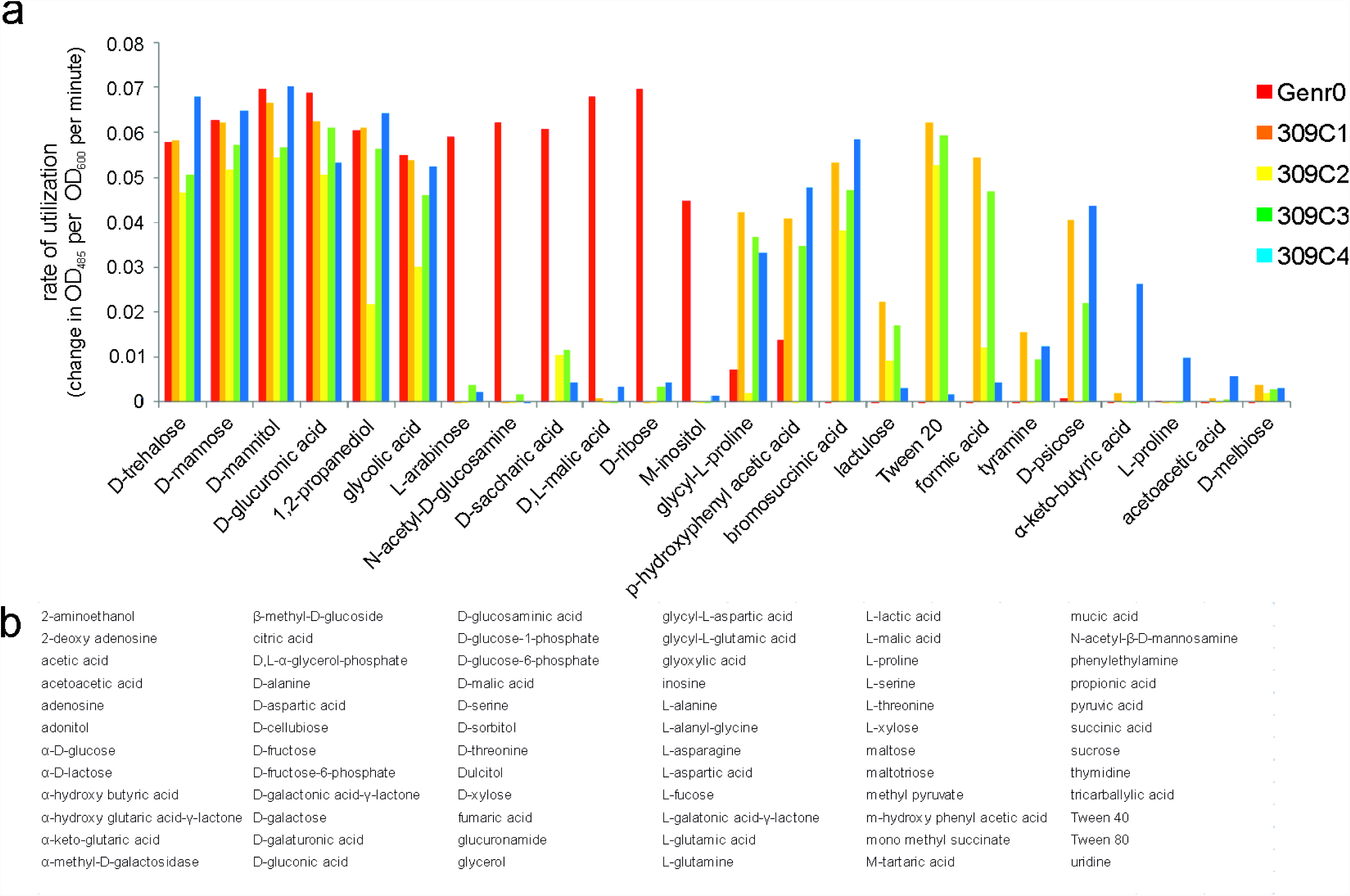
(a) Rate of utilization of carbon sources present in biolog plates. Cells were grown in CDMPC until mid-exponential phase, washed in CDMPC without glucose and resuspended in CDMPC without glucose containing 5 μg/mlchloramphenicol to an OD600 of 2. 150 μl of the culture suspension was added to each of the wells of the biology plate and acid formation was monitored at OD485 for 4 hours. The maximum rates of utilization were calculated and the averages of two separate experiments are shown. Only susbtrates used by at least one strains are shown. (b) Substrates not utilized under the conditions tested.

**Supplementary Figure 5.**
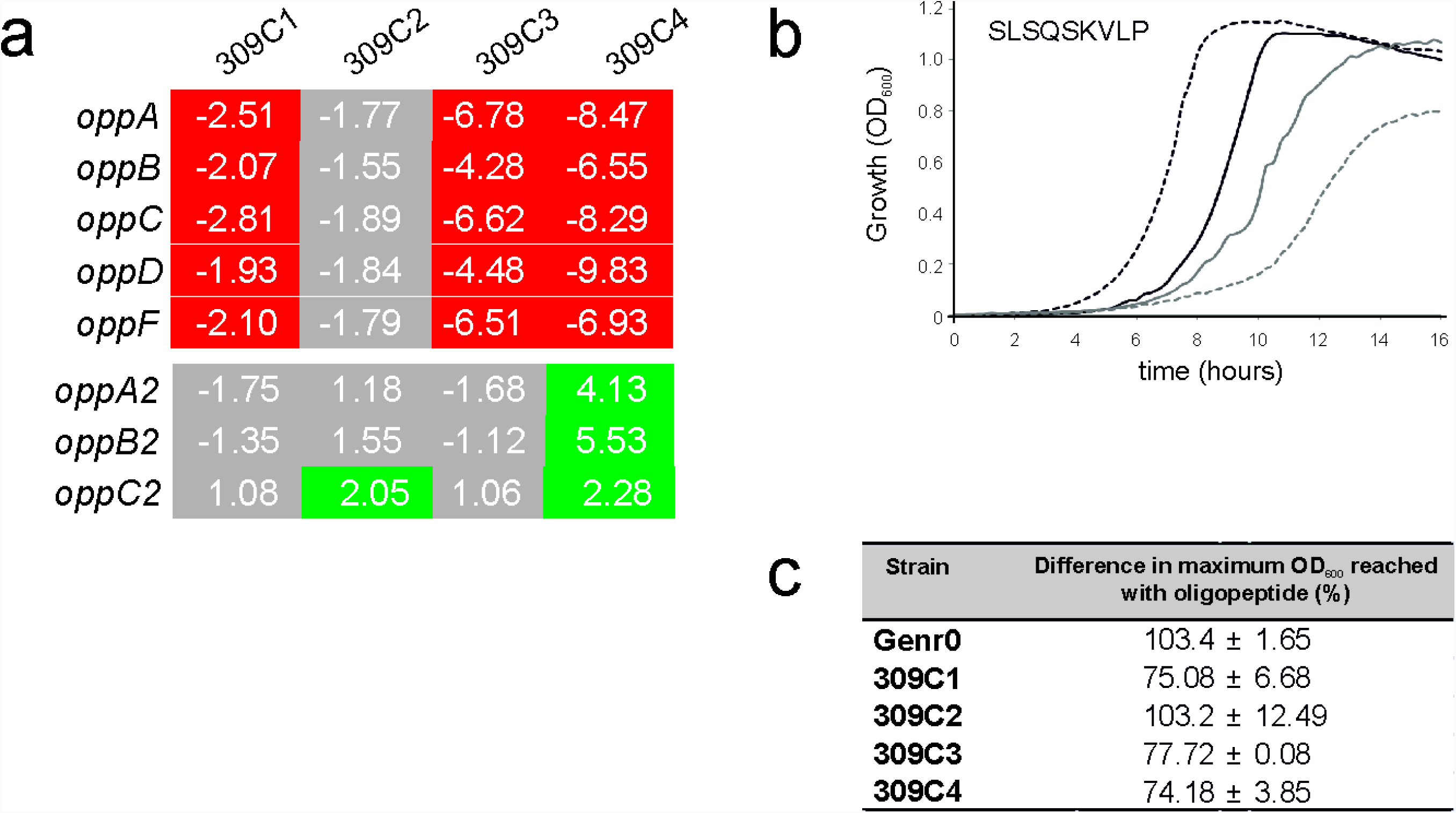
Growth of the evolved strains is inhibited but not abolished when leucine is supplied as a nonapeptide. (a) Differentially expressed genes involved in oligopeptide uptake in *L. lactis* MG1363. Significant changes were considered for genes with a Bayes p-value score ofless than 0.05 and a pfp value of less than 0.05 and are highlighted in green for upregulated genes and in red for down regulated genes. Gene expression changes not meeting the significance cut off values are highlighted in grey. (b) Strains were grown in CDMPC or in CDMPC without leucine but with a leucine-containing nonapeptide. Growth was followed for 16 hours. Shown are Gen0 (black lines) and 309F4 (grey lines) grown in CDMPC (solid lines) or CDMPC with leucine-containing peptide (dashed lines). (c) The difference in the final OD600 values reached after growth for 16 hours are shown for all strains. Shown are the averages between three separate experiments with the standard deviation indicated.

**Supplementary Figure 6.**
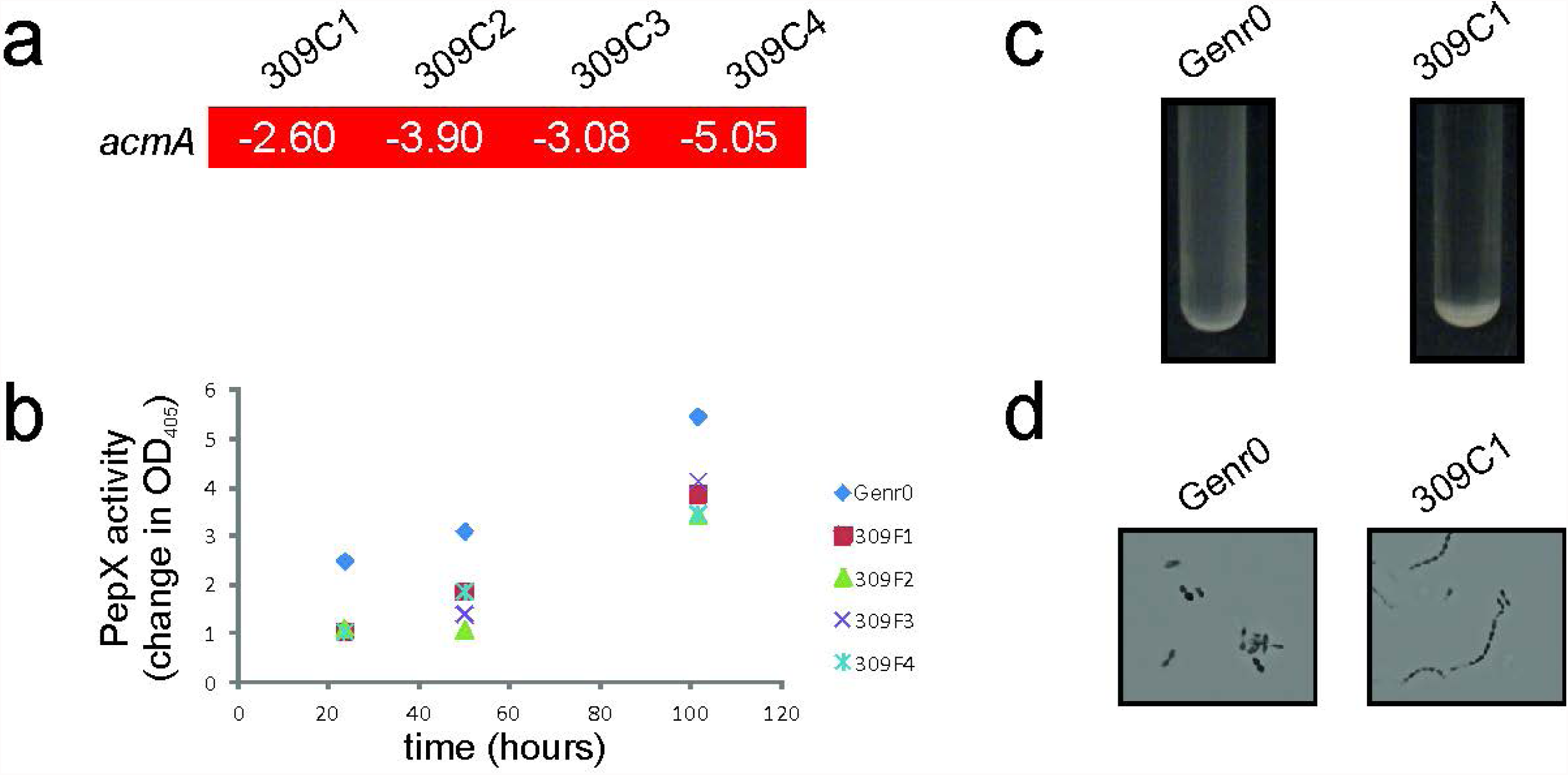
Evolved strains are less prone to lysis on prolonged cultivation than the original strain. (a) Expression changes for acmA are shown. Differentially expressed genes involved in aerobic respiration in L. lactis MG1363. Significant changes were considered for genes with a Bayes p-value score ofless than 0.05 and a pfp value of less than 0.05 and are highlighted in red for down regulated genes. (b) PepX activity was measured in the culture supernatant at the time points indicated. Activity was measured using the chromogenic substrate Ala-Pro-p-nitroanilid and the colour change was monitored at OD405. One representative experiment is shown. (c) Evolved strains sediment during growth. Overnight cultures of Gen0 and 309C1 in CDMPC are shown. (d) Sedimentation is caused by the formation of long cell chains. Typical pictures are shown for Gen0 and 309C1 grown for 16 hours in CDMPC. The cells were visualized using a Zeiss light microscope and a Zeiss digital camera. Magnification, ×1 000.

**Supplementary Figure 7.**
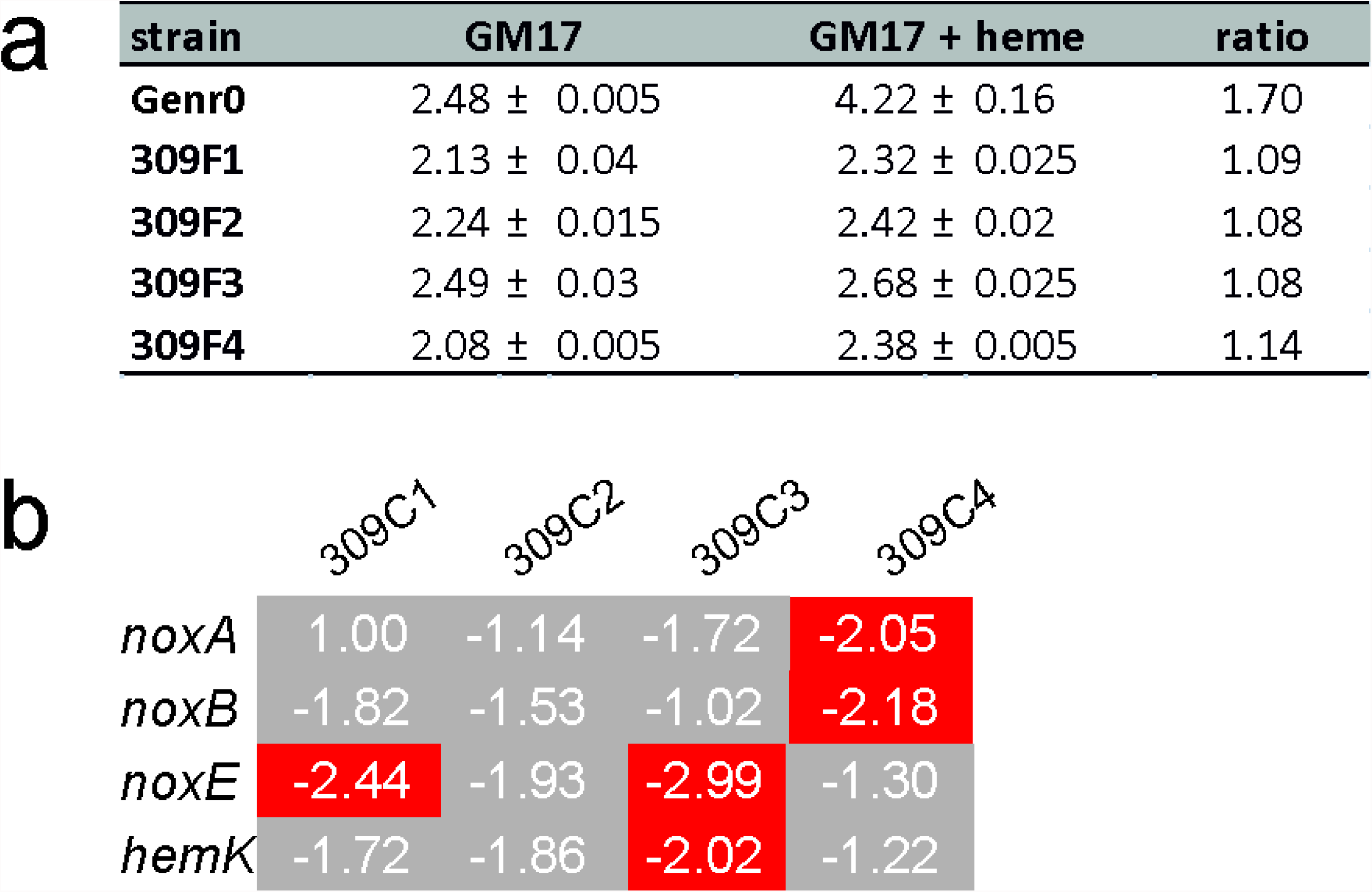
Evolved cells are no longer able to respire when heme is added to the medium. (a) No biomass increase is observed in the evolved strains when heme is added to the growth medium. Strains were grown in GM17 medium for 16 hours under aerobic conditions with and without heme which was added to 2 μg/ml. Shown are the average values from three separate experiments with the standard deviation indicated. (b) Differentially expressed genes involved in aerobic respiration in *L. lactis* MG1363. Significant changes were considered for genes with a Bayes p-value score of less than 0.05 and a pfp value of less than 0.05 and are highlighted in red for down regulated genes. Gene expression changes not meeting the significance cut off values are highlighted in grey.

**Supplementary Figure 8.**
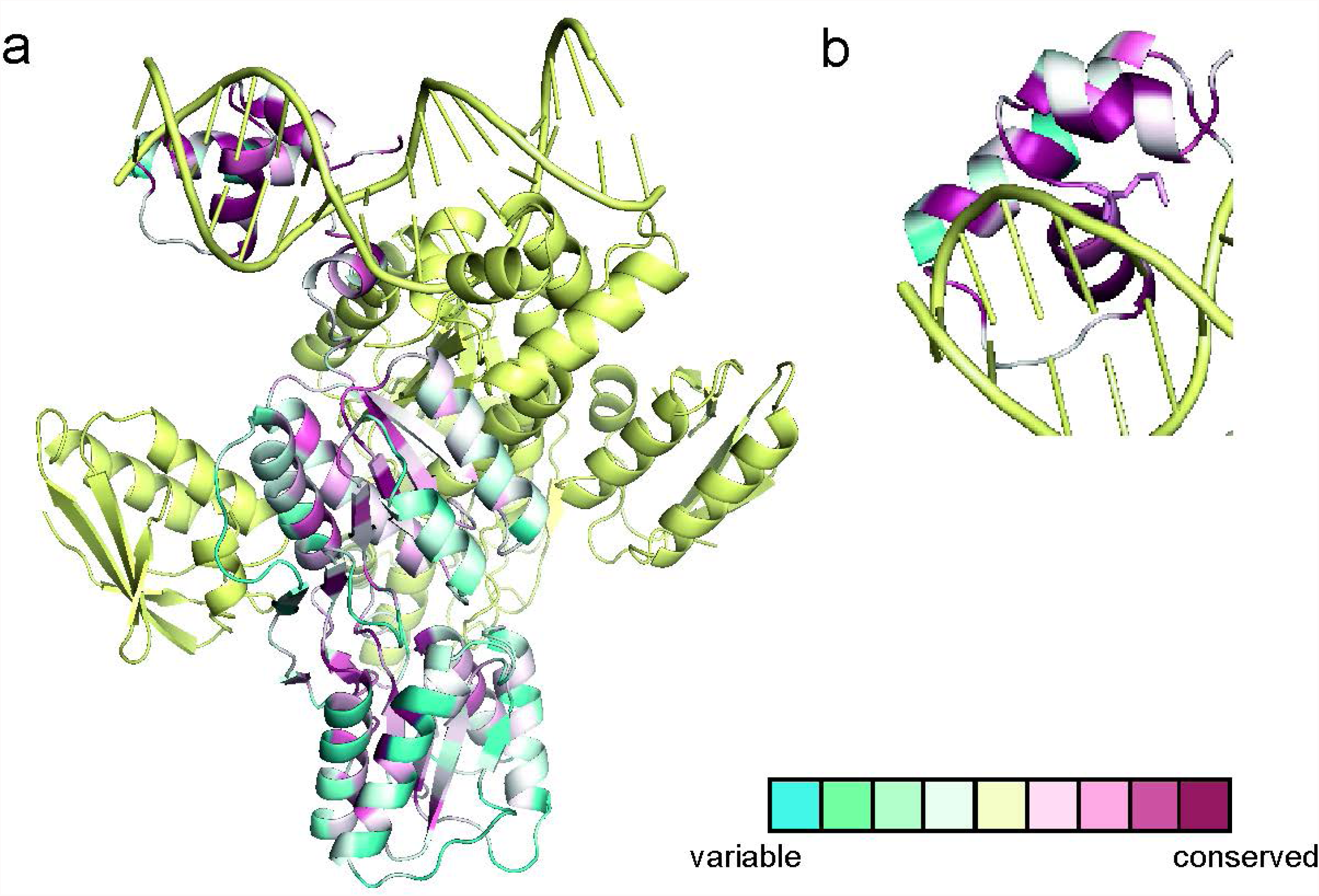
The DNA binding domain of CcpA is highly conserved. (a) Using consurf db (Ashkenazy et al. 2010) and the *B. subtilis* CcpA-HPr complex bound to a synthetic cre site (3OQN) as a starting structure, the level of amino acid sequence conservation among LacI transcriptional regulators was analysed. One molecule of CcpA is colored according to the level of conservation within the LacI family of transcriptional regulators, HPr and the DNA are colored yellow. Dark purple indicates 100% conservation while blue indicates extensive sequence variation. (b) View of the DNA-binding domain with the side chain of Met19 (numbering according to *L. lactis* MG1363) is shown.

**Supplementary Figure 9.**
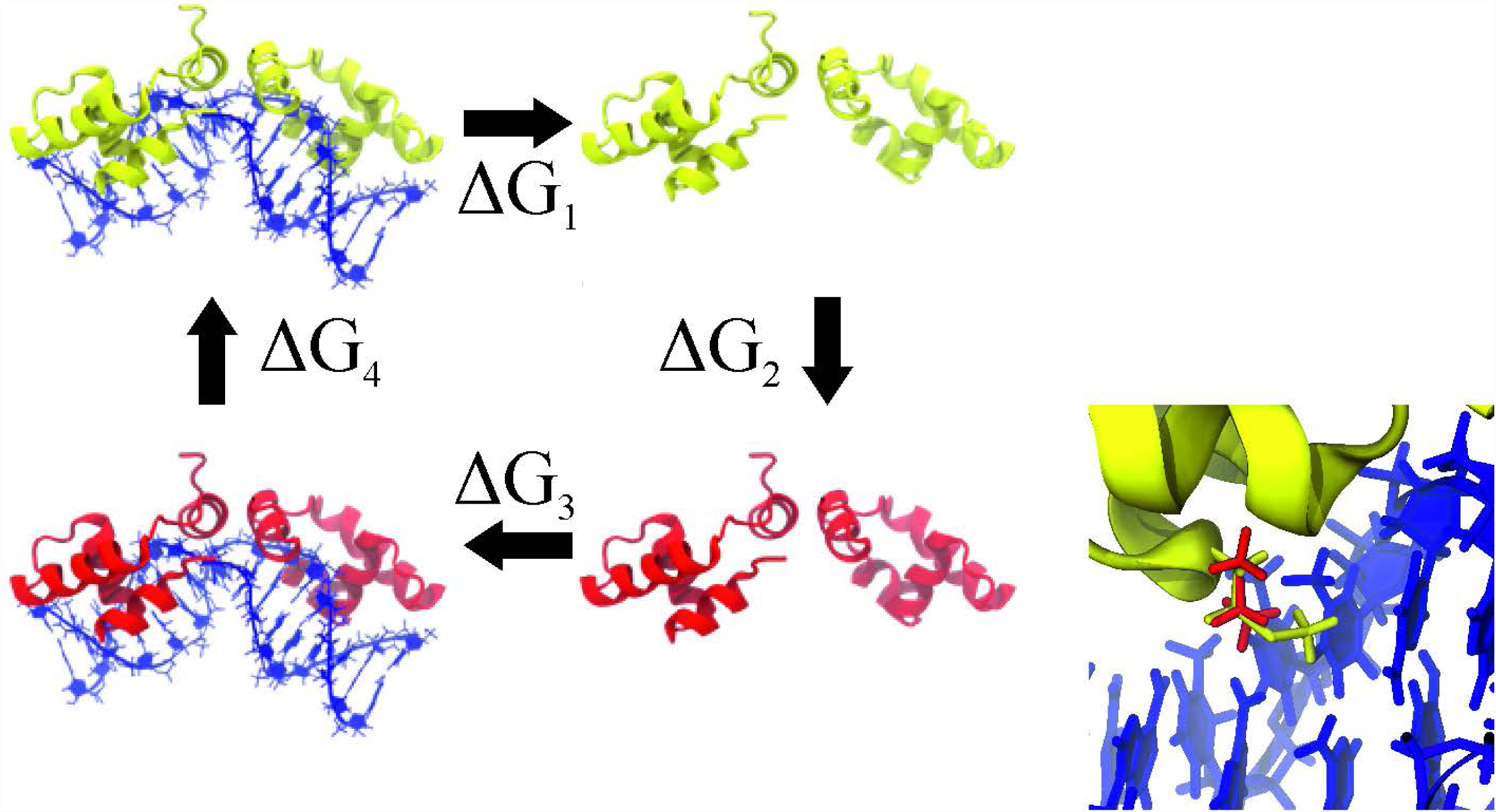
The thermodynamic cycle to calculate the relative binding free energies. The wild type protein is shown in yellow and the mutated protein in red. Note that while the color of the whole protein is different only a single residue is changed in both monomers of the mutated protein. The relative binding free energy is ΔG1 + ΔG3 which, based on the cycle, is equal to -(ΔG2 + ΔG4). The inset highlights the DNA binding domain of CcpA, with the methionine-19 position in red, and isoleucine at the same position in yellow.

**Supplementary Figure 10.**
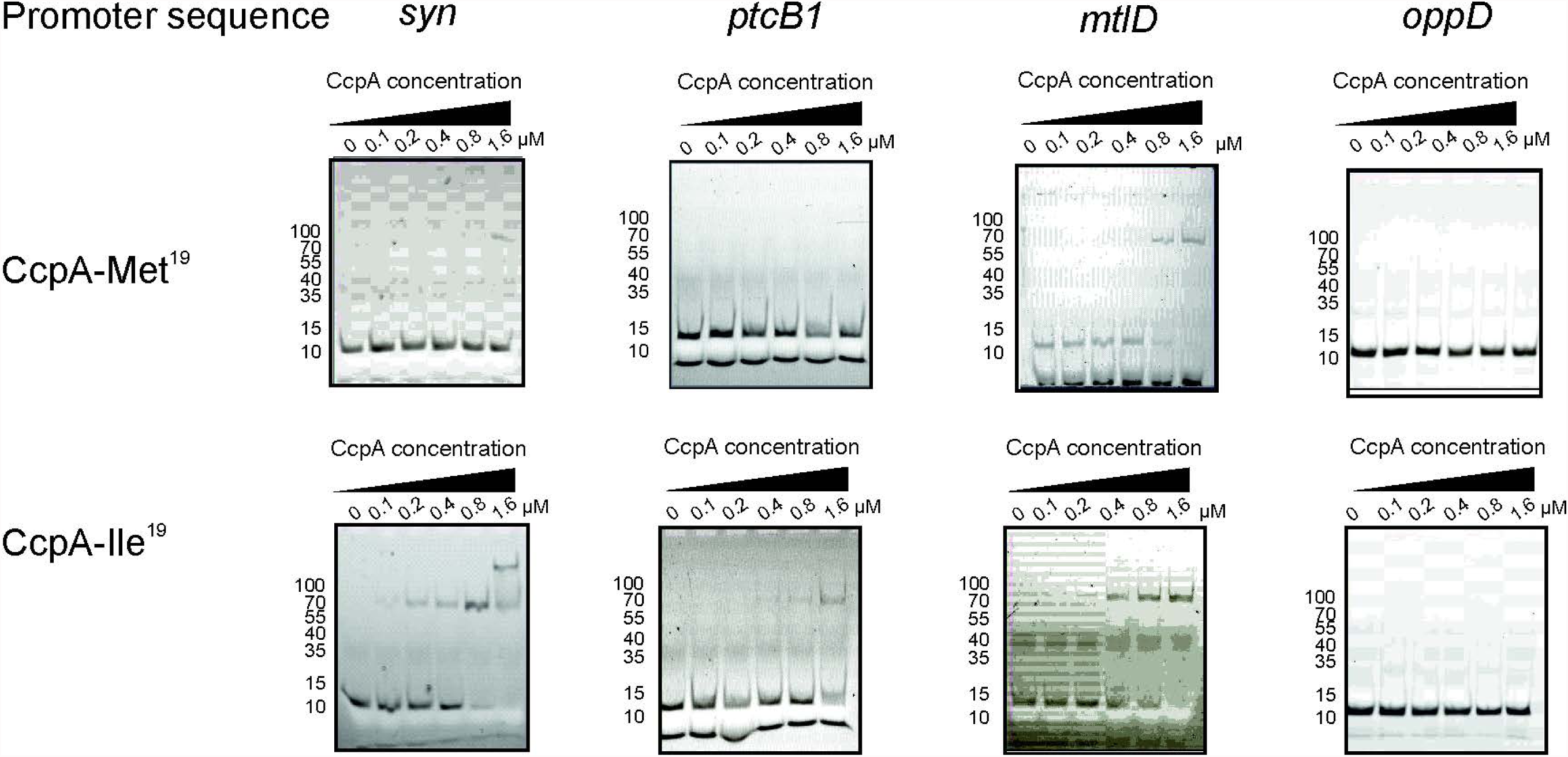
Binding of CcpA to cre sites in vitro. The binding of CcpA-Met19 and CppA-Ile19 was tested with DNA sequences identified as cre sites upstream of *ptcB* and *mtlD* as well as a perfect cre site referred to as syn. As a control the CodY recognition site upstream of *oppD* was also tested.

**Supplementary Figure 11.**
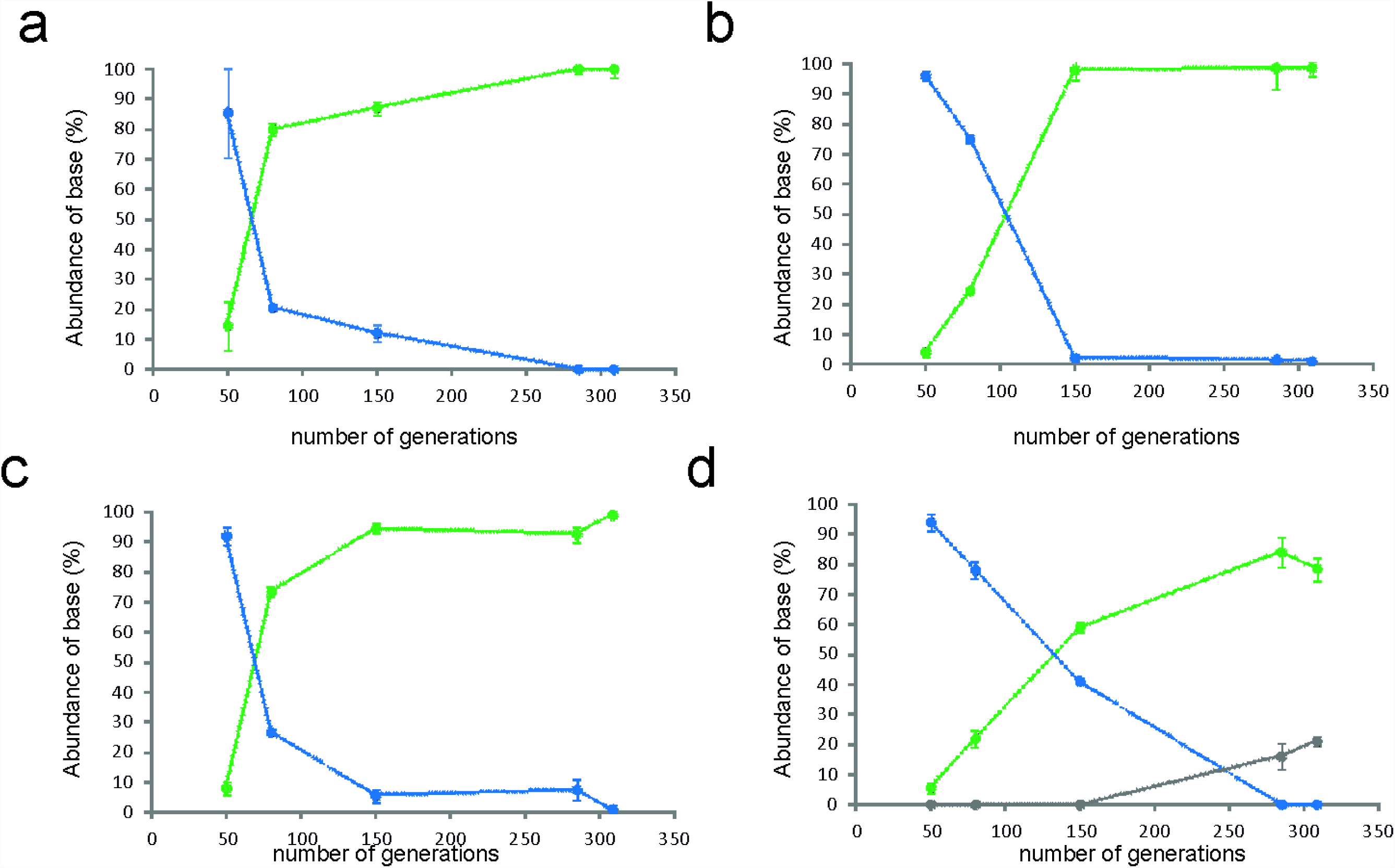
*ccpA* accumulates the Met19 to Ile early in evolution experiment and quickly takes over the population. The abundance of the three CcpA species is shown. Indicated are the averages of two pyrosequencing reactions and in the case of strain 309C4, the abundance of the Thr mutation was verified by Sanger sequencing. The PCR was performed from the frozen stocks sampled during the continuous culture process. In green is indicated the prevalence of the Ile mutant, in blue wild type CcpA and in grey the Thr mutant. (a) 309C1, (b) 309C2, (c) 309C3, (d) 309C4.

**Supplementary Figure 12.**
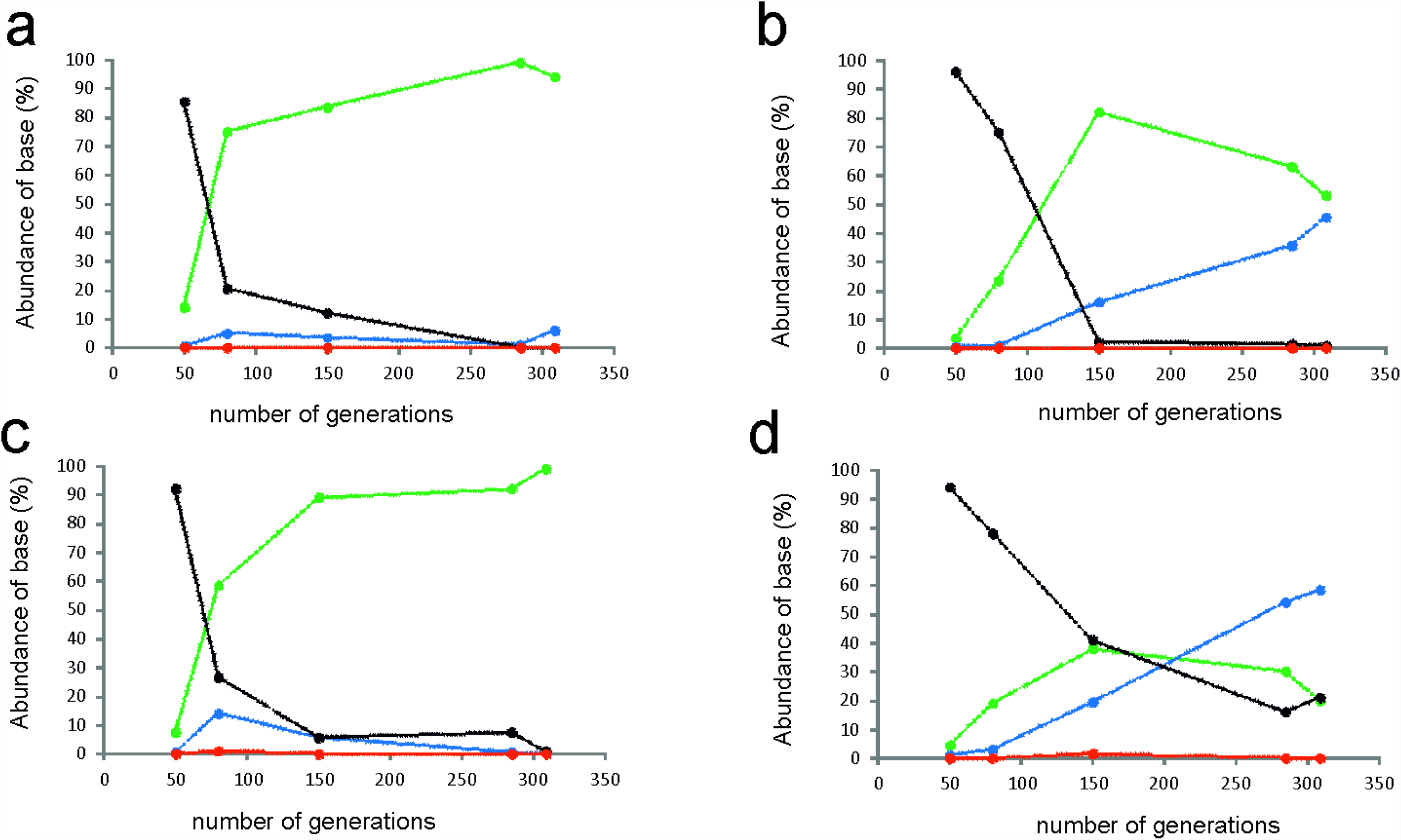
Abundance of nucleotides at third position in the Met19 codon. Indicated are the averages of two pyrosequencing reactions. The PCR was performed from the frozen stocks sampled during the continuous culture process. In green is indicated the prevalence of adenosine, in red thymine, in blue cytosine and in black guanine. The average standard deviation between the separate reactions was 4.4. (a) 309C1, (b) 309C2, (c) 309C3, (d) 309C4.

**Supplementary Figure 13.**
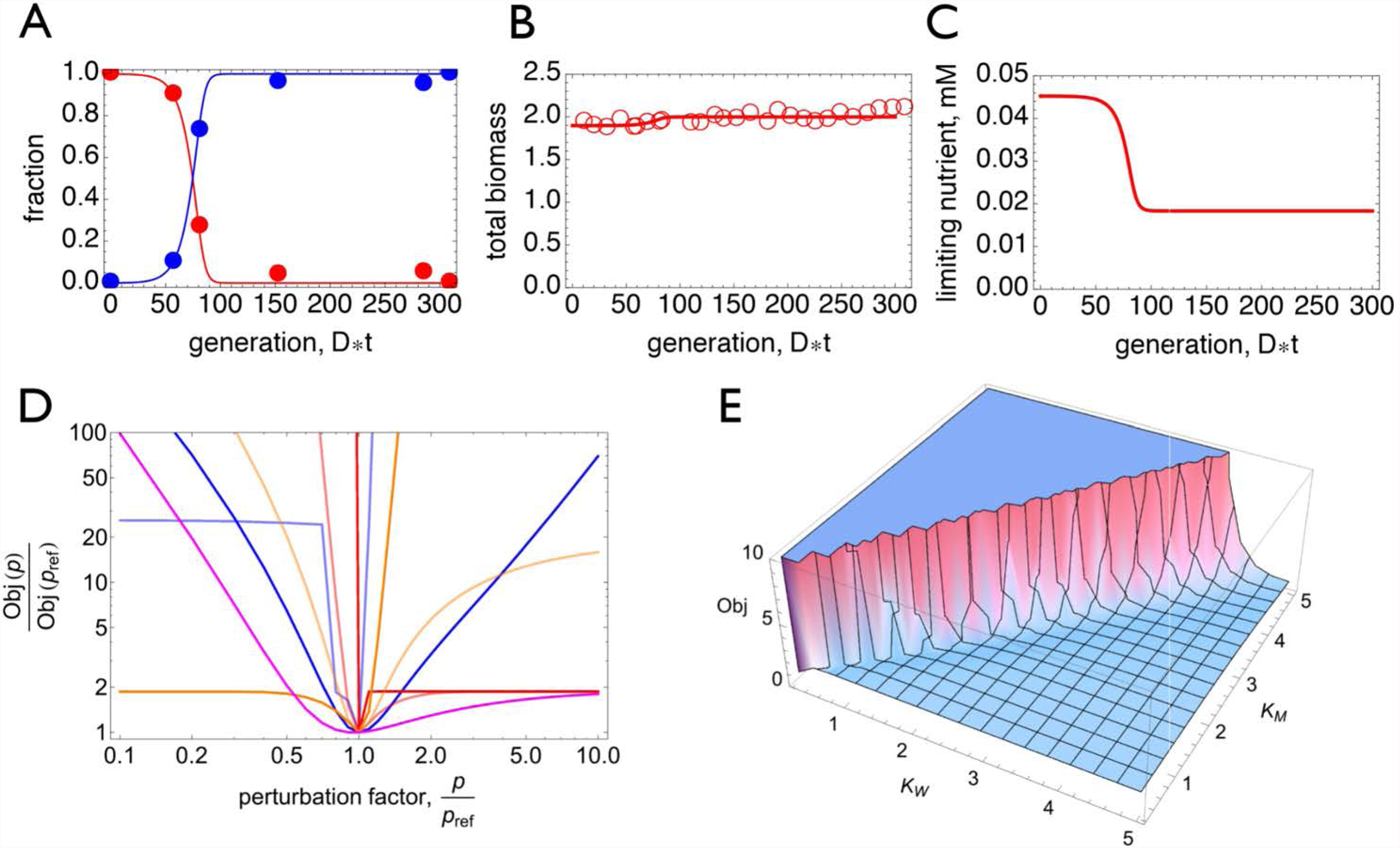
Overview of the fitting results. A & B. The fitted model (eq. 2 and 3) and the experimental data. C. Predicted dynamics of the concentration of the limiting nutrient in the chemostat. The mutant wins the competition with the wild type because it can grow at a specific growth rate equal to D at a lower limiting nutrient concentration than the wild type. D. Influence of parameter values on the objective, indicating that all the parameter values reach their optimal values (reported in Supplementary Table 1) when the fit objective reaches its minimal value. E. Illustration that the Monod constant of the wild type and the mutant cannot be independently estimated; i.e. the same objective value is obtained for different values of those parameters. The plot suggests that the ratio of Kw/Km can be fitted but not their individual values.

**Supplementary Table 7:**
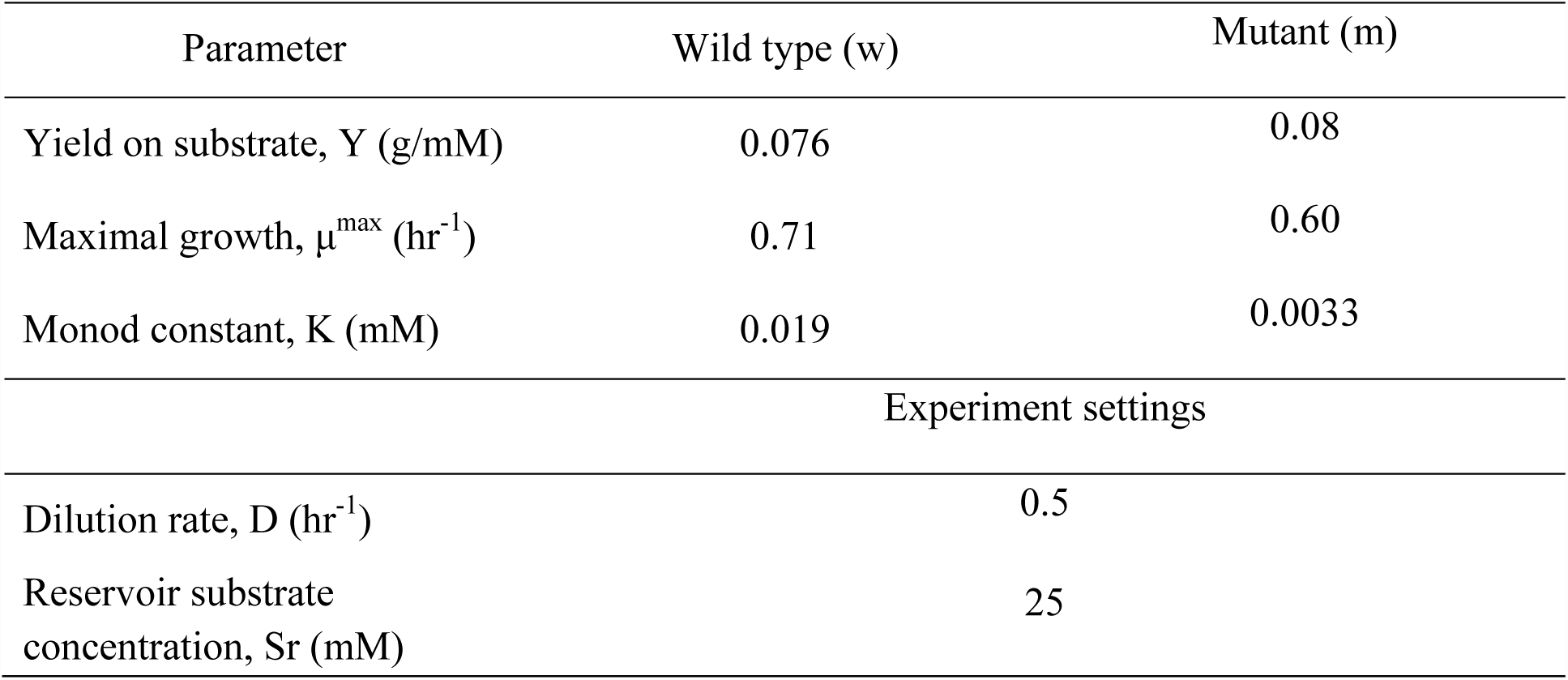
Overview of parameters (fitted and experimental settings).

